# The impact of bottlenecks and inbreeding on the genome of the endangered Pyrenean desman

**DOI:** 10.1101/2020.07.25.199281

**Authors:** Lídia Escoda, Jose Castresana

**Author notes:** Corresponding author: Jose Castresana.

## Abstract

The Pyrenean desman (*Galemys pyrenaicus*) is a small semiaquatic mammal endemic to the Iberian Peninsula. Despite its limited range, this species presents a strong genetic structure due to past isolation in glacial refugia and subsequent bottlenecks. Additionally, some populations are highly fragmented today as a consequence of river barriers, causing substantial levels of inbreeding. These features make the Pyrenean desman a unique model in which to study the genomic footprints of differentiation, bottlenecks and extreme isolation in an endangered species. The complete genome of the Pyrenean desman was assembled using a Bloom filter-based approach. An analysis of the 1.83 Gb reference genome and the sequencing of five additional individuals from different evolutionary units allowed us to detect its main genomic characteristics. The population differentiation of the species was reflected in highly distinctive demographic trajectories. A severe population bottleneck during the postglacial recolonization of the eastern Pyrenees created the lowest genomic heterozygosity ever recorded in a mammal. Moreover, isolation and inbreeding gave rise to a high proportion of runs of homozygosity (ROH). Despite these extremely low levels of genetic diversity, two key multigene families from an eco-evolutionary perspective that need to be genetically variable, the major histocompatibility complex and olfactory receptor genes, showed heterozygosity excess in the majority of individuals. Furthermore, these two classes of genes were significantly less abundant than expected within ROH. These results allow us to characterize important genomic health indicators for each individual, information that may be crucial for the conservation and management of the species.

## Introduction

Complete genomes of endangered species are helping to identify features of individuals and populations that can be critical for *in situ* and *ex situ* conservation (Xue et al. 2015; Abascal et al. 2016; Benazzo et al. 2017; Ekblom et al. 2018; Westbury et al. 2018; Saremi et al. 2019). To adequately manage species, it is essential to know not only which populations may be most threatened, but also which individuals from healthy populations may be optimal for genetic rescue or captive breeding. One of the important characteristics that must be considered in conservation is genetic diversity, which can be measured in individuals as the proportion of heterozygous positions in the genome. The heterozygosity rate or SNP density has been shown to vary greatly between different species (Prado-Martinez et al. 2013). When considering only mammalian species of conservation concern, this value can be as low as 14 heterozygous positions or SNPs per million bases (SNPs/Mb) in a population of foxes on a small oceanic island (Robinson et al. 2016) and can reach as many as 1,200 SNPs/Mb in some orangutan populations (Locke et al. 2011), thus extending over two orders of magnitude. Low heterozygosity is generally caused either by population bottlenecks that have occurred in recent evolutionary history or current population declines, and it is unclear as to whether there is a critical value below which an individual or population can be considered at risk.

More important than the average heterozygosity rate is the variability of genetic diversity in the genome. The strongest heterozygosity fluctuations on chromosomes are caused by inbreeding. If inherited from a recent common ancestor, both copies of some chromosome blocks can be identical in inbred individuals, forming the so-called runs of homozygosity (ROH) (Ceballos et al. 2018). The proportion of the genome in ROH is the inbreeding coefficient. In many species, including humans, inbreeding leads to reduced fitness due to the presence of lethal mutations in homozygosis. When inbreeding is widespread and there is a positive correlation between individual inbreeding coefficients and fitness in a population, inbreeding depression may occur, often leading to an extinction vortex in the short term (Kardos et al. 2016). Nevertheless, this pernicious association is not seen in populations in which lethal mutations have been purged during bottlenecks in their recent population history (Keller and Waller 2002). In any case, knowing the inbreeding coefficient of individuals is critical for managing populations.

Proteins that directly interact with the environment are particularly interesting in the context of genetic diversity as these require a high degree of inter- and intra-locus variability to function properly. Two protein families are key in this regard: the major histocompatibility complex (MHC), which is one of the principal components of the immune system (Abduriyim et al. 2019; Radwan et al. 2020); and the olfactory receptors (OR), which involves the largest multigene family in mammals (Hughes et al. 2018). Due to balancing selection, a huge diversity of alleles is found in some MHC genes, consequently raising heterozygosity in these particular loci (Vandiedonck and Knight 2009). OR loci are also suspected to be more heterozygous due to the advantage conferred by this state (Alonso et al. 2008). However, it is not yet known whether certain heterozygosity levels can be maintained in these genes when populations have extremely low genetic diversity or significant inbreeding levels. Another interesting question is whether ROH regions in inbred individuals can include genes for which heterozygosity is an advantage or if these genes are less abundant in ROH.

The Pyrenean desman (*Galemys pyrenaicus*) is one of the only two extant species of the mammalian subfamily Desmaninae, a lineage that was composed of a large number of species during the Neogene (McKenna et al. 1997), and therefore has a high extinction rate. This small mammal is endemic to the Iberian Peninsula. It lives in small rivers with clean waters, and possesses major adaptations for semi-aquatic life (Palmeirim and Hoffmann 1983; Kryštufek and Motokawa 2018). Due to its shrinking distribution, it is classified as vulnerable in the IUCN Red List and some of its populations are highly threatened (Fernandes et al. 2008). It has been shown that its genetic structure is very strong, being subdivided into five populations (evolutionarily significant units), likely to be of glacial origin (Igea et al. 2013; Querejeta et al. 2016). A ddRAD-based study on the species revealed extremely low heterozygosity in some individuals from the eastern Pyrenees, probably as a consequence of repeated bottlenecks during the postglacial recolonization of these mountains (Querejeta et al. 2016). In addition, it has been demonstrated that the isolation of populations in the upper parts of rivers due to the construction of artificial barriers, such as dams, is leading to extremely high inbreeding levels in some areas (Escoda et al. 2017; Escoda et al. 2019). Having the complete genome of the Pyrenean desman would allow us to study how the combination of strong population bottlenecks and high inbreeding levels is reflected in the genomic landscape of a threatened species. No genome from this species has been obtained so far and the closest sequenced genome comes from a mole of a different subfamily. Here, we provide the first draft genome assembly and annotation of the Pyrenean desman and reveal highly peculiar patterns of genomic diversity in the whole genome as well as in the MHC and OR genes.

## Results

### Pyrenean desman samples sequenced

A total of six Pyrenean desman individuals covering the majority of the species distribution range (Figure 1, supplementary Table S1) was sequenced using Illumina libraries of various sizes. To assemble the reference genome, genomic DNA from a male from the eastern Pyrenees was sequenced at 121x coverage (Table S2) while five additional desmans from other locations (western Pyrenees, northwest and southeast Iberian Range, Central System, and West of the Iberian Peninsula, i.e., four out the five main populations) were resequenced (Figure 1, Table S3).

**Figure 1.**
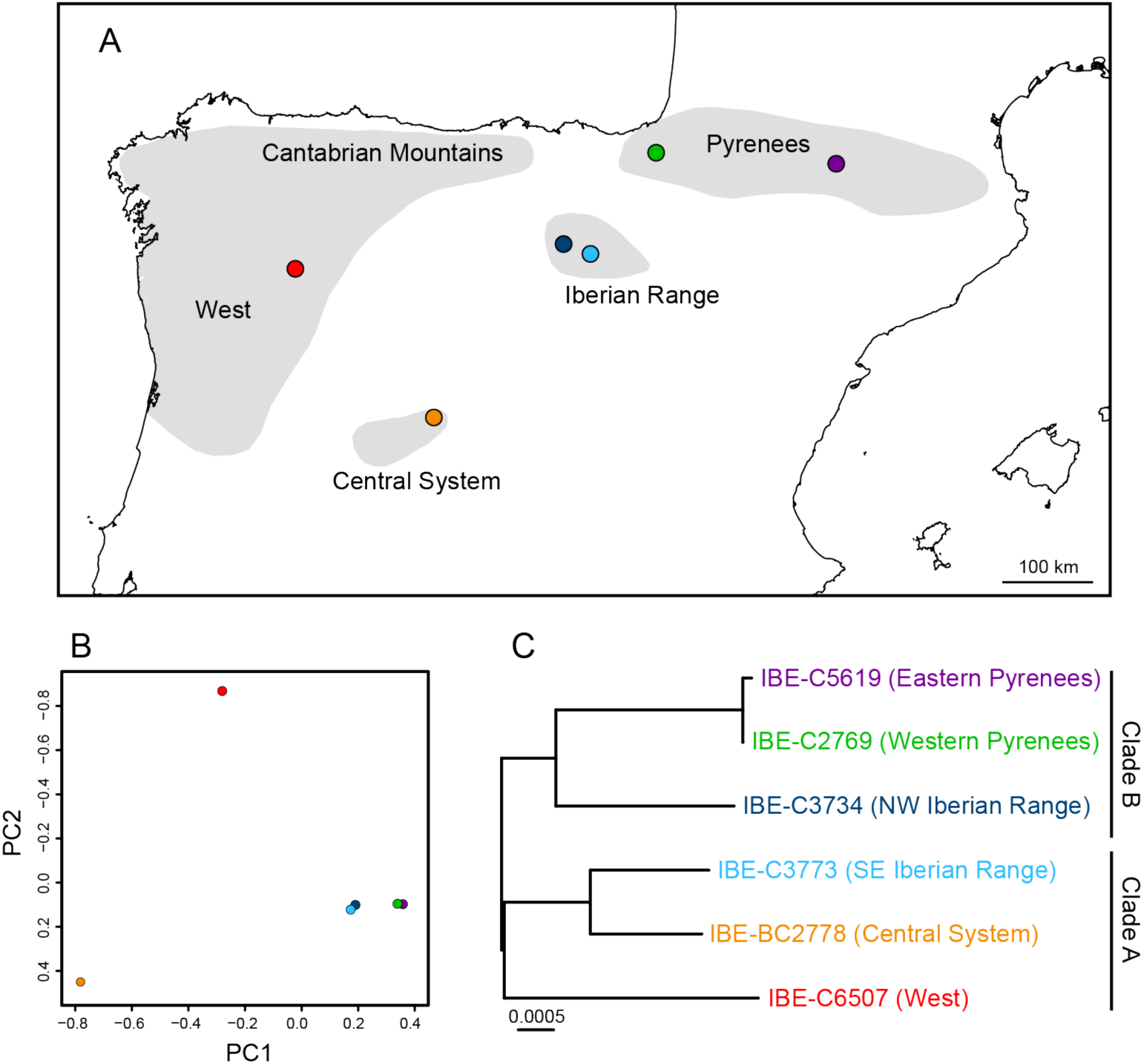
(A) Map of the Iberian Peninsula showing the distribution of the Pyrenean desman (shadowed areas) and the locations of the sequenced individuals. (B) Principal component analysis of the genotypes. (C) Maximum-likelihood phylogenetic tree of the mitochondrial genomes, in which the main clades are indicated. The root of the tree was placed between clades A and B and the scale is in substitutions/site.

### Genome assembly and Bloom filter parameterization

To assemble the *de novo* genome, we used a strategy with relatively low computational memory requirements based on the ABySS program with the Bloom filter option (Jackman et al. 2017). Since this method has not been thoroughly tested for large genomes, we first searched for the optimal parameters of the Bloom filter to assemble the desman genome. A total of 32 combinations of Bloom filter parameters, k-mer values, and other ABySS options were tested (Table S4). The summary statistics used indicated that very different results can be obtained depending on the parameters. The parameters that gave the best overall results were a minimum k-mer count threshold for the Bloom filter (kc) of 2, a Bloom filter size (B) of 80G, and a number of Bloom filter hash functions (H) of 4, together with a k-mer size (k) of 100 and other ABySS parameters detailed in Table S4. The 1.83 Gb final assembly had a scaffold N50 of 8.5 Mb (Table S5) and 96.3% of the mammalian BUSCO core genes (Table S6). Although assemblies with higher N50 values were obtained, these involved a larger number of scaffolds, more gaps, or lower numbers of BUSCO genes (Table S4). We further evaluated the accuracy of the desman genome assembly by mapping the short-insert sequencing data to the assembled genome: 82.5% of the reads were mapped and a low-coverage peak corresponding to the Y- and X-chromosomes was observed, as expected for a male (supplementary Figure S1A). The GC content was 41.7% and showed a variability across chromosome regions typical of a mammalian genome (Figure S2).

The high-coverage sequencing data we obtained did not allow us to use the standard ABySS mode for comparison, due to the large amount of RAM memory required. A comparison with an already assembled dromedary genome using the standard ABySS mode (Fitak et al. 2016) and a new Bloom filter assembly showed a better scaffold N50 and a greater longest sequence for the Bloom filter assembly, although the BUSCO statistics were slightly better for the standard assembly (Table S7). The alignment of the two assemblies showed that they were very similar (Figure S3).

### Gene prediction

After detecting repetitive elements (Table S8) and masking the genome, we predicted 20,936 protein-coding genes using the MAKER2 pipeline (Holt and Yandell 2011). The annotation edit distance (AED) of the genes, which provides a measure of the prediction congruence, showed that 95% of the genes have a score lower than 0.5 (Figure S1B), indicating a well-annotated genome (Campbell et al. 2014). Other features of the predicted genes that indicated that the Bloom filter-based assembly of the Pyrenean desman is equivalent to any other properly assembled mammalian genome include a bimodal distribution of intron length (Figure S1C), as observed in other mammalian genomes (Piovesan et al. 2016), and some genes with coding sequences (CDS) longer than 100,000 bp (Figure S1D), such as titin (Labeit and Kolmerer 1995).

The gene sequences of two important multigene families were retrieved and aligned. The phylogenetic trees of 26 class-I MHC α chain (MHC-I-α) (Figure S4A) and 529 OR (Figure S4B) genes together with those of other mammals indicated a large diversity of genes in both families.

### Genetic structure and demographic history

The principal component analysis (PCA) of the genotypes agreed, in general terms, with the geographic proximity of the individuals (Figure 1B). On the other hand, the maximum-likelihood phylogenetic tree of the assembled mitochondrial genomes (Figure 1C) showed an important mito-nuclear discordance for the individual from the SE Iberian Range, whose geographic and nuclear proximity to the other individual from the Iberian Range is not reflected in the mitochondrial tree, corroborating previous work (Igea et al. 2013; Querejeta et al. 2016; Escoda et al. 2017).

The pairwise sequentially Markovian coalescent (PSMC) analysis (Li and Durbin 2011) revealed that all the populations experienced a general decline together with substantial fluctuations in their effective sizes during the time covered by the plot, of which the last ∼300 thousand years showed the best resolution (Figures 2 and S5). When compared with the major climatic events occurred in this time interval (Clark et al. 2009; Dahl-Jensen et al. 2013), the two population size peaks observed approximately coincide with the beginning of the two interglacial periods of this time (Eemian and Holocene). Within this general pattern, there were important differences among individuals. The demographic fluctuation patterns were similar for the two desmans from the Iberian Range and, to a certain extent, the one from the western Pyrenees. The individual from the West of the Iberian Peninsula showed a delayed decline and the highest current effective population size. The desman from the Central System presented a high population size peak during the Eemian interglacial and a large decline since then. Finally, the curve of the desman from the eastern Pyrenees revealed an extremely small effective population size and its data only covered a short period of time, probably due to its exceptionally low heterozygosity (see below).

**Figure 2.**
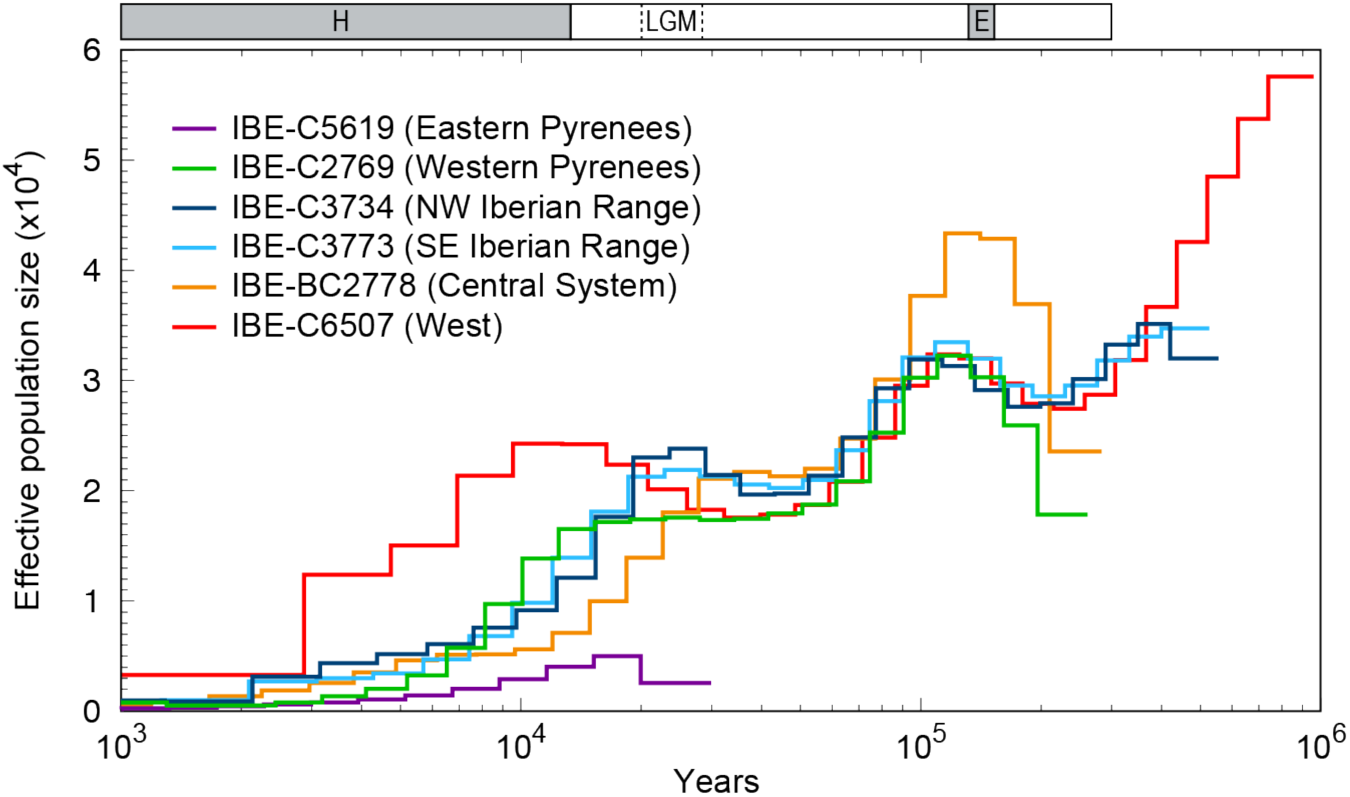
Historical effective population size of the Pyrenean desman individuals, inferred by PSMC. The result is scaled with a mutation rate of 5 × 10^−9^ mutations/site/generation and an average generation time of 2 years. The last two interglacial periods, Holocene (H) and Eemian (E), are indicated with grey boxes and the Last Glacial Maximum (LGM) with dashed lines.

### Genomic heterozygosity and runs of homozygosity (ROH)

The autosomal genome heterozygosity rate varied greatly between the six desmans and is among the lowest found in mammals (Table S9; Figure S6). It ranged between 12 and 463 SNPs/Mb, with an average of 202 for all individuals. Similar values were obtained with less strict minimum depth of coverage filtering, which allowed for a larger SNP count (average of 189 SNPs/Mb for all individuals; Table S9). The desman from the eastern Pyrenees, with 12 SNPs/Mb (result obtained with both filtering options), has the lowest heterozygosity recorded in a mammal so far (Figure S6). In comparison with the ddRAD data available for three of the desmans, the ddRAD heterozygosity values were lower than those from the genome data: an average of 236 SNPs/Mb for ddRAD data versus 321 or 300 (depending on the minimum depth of coverage) for the genomes of the three individuals (Table S9). This ∼25% underestimation is probably due to the fact that ddRAD does not capture the most divergent genome sequences. However, the comparative value of heterozygosity rates estimated with ddRAD remains valid, as seen in other works (Wright et al. 2020).

When we calculated the heterozygosity in 100 kb windows and plotted the values across the scaffolds, we found that most of the desmans presented very long ROH, with important variations in lengths and patterns among individuals (Figure 3). To calculate the proportion of ROH for the genome of each individual, which is equivalent to its inbreeding coefficient, we used four different approaches. Two of the methods (PLINK and BCFtools/RoH) are based on the variable positions across all the individuals, while the other two (ROHan and the simple proportion of 100 kb windows with 0 heterozygous positions) use the complete information from each individual genome, independently of the variants found in other genomes. The values of the four methods were similar and highly correlated (Table S10). They were also congruent with the inbreeding coefficients estimated from populational ddRAD data available for three of the desmans. These results indicate that methods for estimating ROH independently of the population background can be perfectly valid when no population data is available. They also show that all the desmans had very high values of the inbreeding coefficient. For example, when calculated as the proportion of homozygous 100-kb windows (a simple method giving an average that is very close to the ddRAD data), these values varied between 0.10 for the individual from the West of the Iberian Peninsula and 0.69 for the individual from the eastern Pyrenees (Table S10; Figure 3).

**Figure 3.**
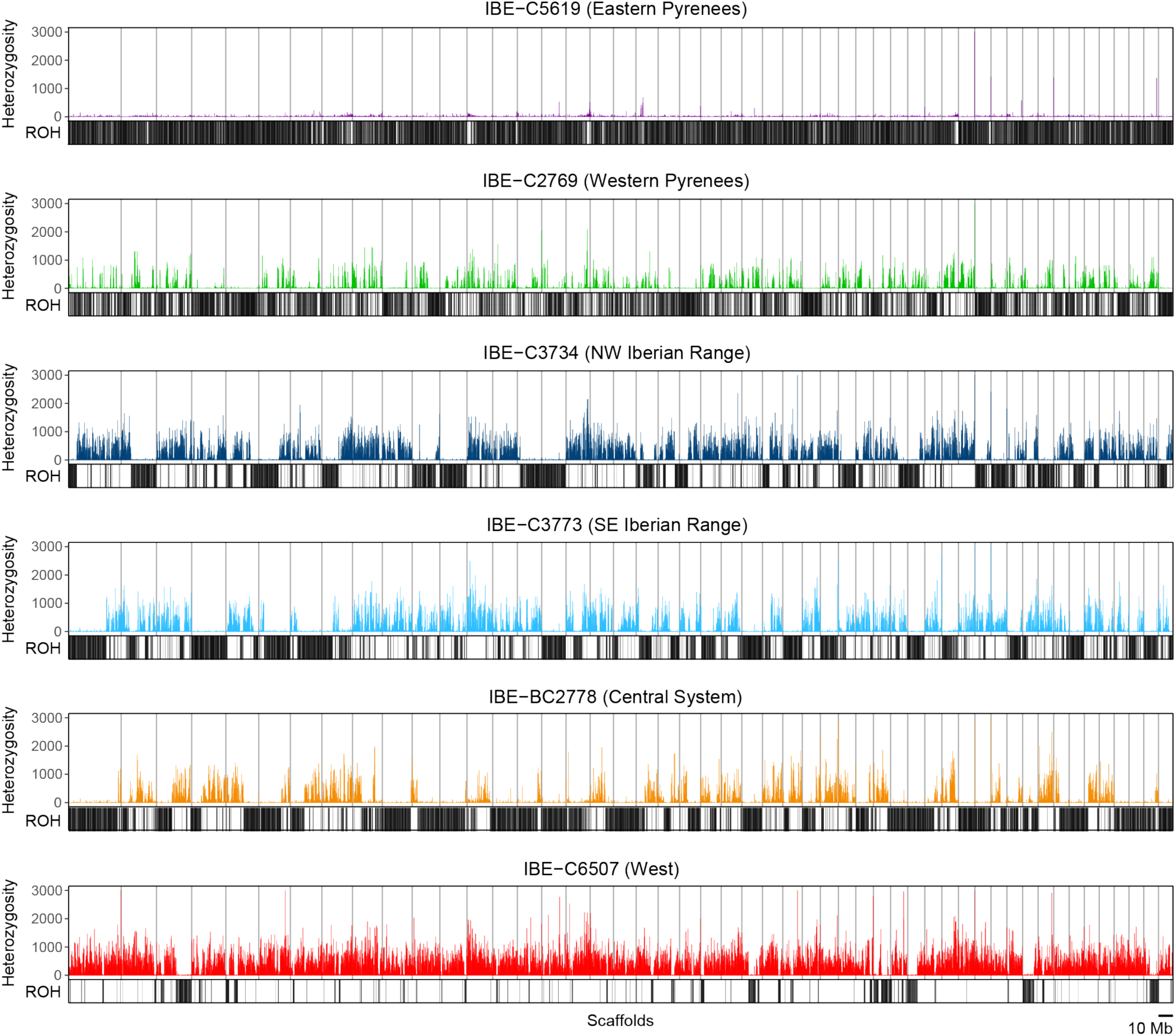
Heterozygosity in SNPs/Mb of the scaffolds longer than 10 Mb for each Pyrenean desman specimen. The black vertical lines under each graph indicate 100-kb windows with 0 SNPs used to calculate the proportion of ROH. Outlier windows with >3,000 SNPs/Mb (between 1 to 4 per individual) were truncated for visual purposes.

### Heterozygosity excess and ROH deficiency in proteins that need to be variable

The low genomic heterozygosity observed in the whole genome is also reflected in the exons (mean of 171 SNPs/Mb across all exons and individuals; Table S11 and Figure 4A). A low genetic diversity may affect the adequate functioning of genes that need to be variable, such as the MHC or OR genes. To understand how this extreme reduction in genetic variability affected these particular genes, we calculated the heterozygosity in their exons (Table S11 and Figure 4A). For the MHC-I-α exons, most individuals presented much higher heterozygosity values (2,052 SNPs/Mb, i.e., 10.9 times higher on average than the whole genome). The desman from the West of the Iberian Peninsula showed the highest heterozygosity (16.9x excess) whereas the desmans from the western Pyrenees and the Central System showed a very low excess. OR exons also presented heterozygosity excess with respect to all exons (605 SNPs/Mb on average, representing a 3.2x excess). In this case, the heterozygosity excess was more similar between all the individuals.

**Figure 4.**
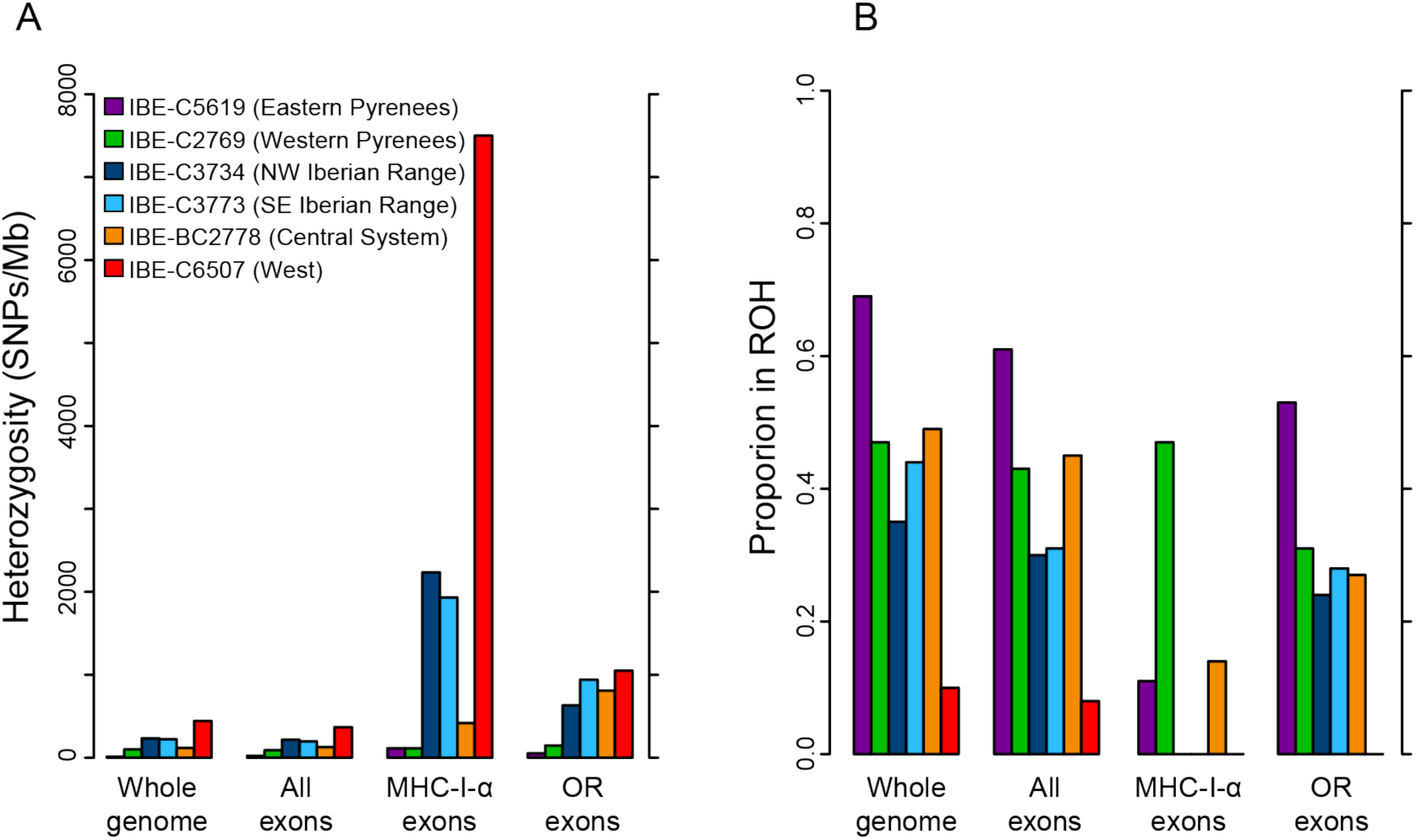
(A) Heterozygosity rate in exons of MHC-I-α and OR of the sequenced Pyrenean desmans, given in SNPs/Mb, in comparison with the heterozygosity of the whole genome and all the exons. (B) Proportion in ROH regions of exons of MHC-I-α and OR in comparison with the ROH proportion calculated for the whole genome and all the exons (colors are as in panel A).

Another important question is whether genes that need to be variable are contained or not within ROH segments. If the presence of such genes in ROH is suboptimal, a lower proportion of their exons is expected in ROH. The expected value is the proportion of homozygous 100-kb windows from the whole genome, in other words, the inbreeding coefficient (Table S10). When calculated for the entire set of exons, this proportion was 1.1 times smaller, on average, than that expected (0.37 versus 0.41; Table S12 and Figure 4B). Despite this small difference, it was highly significant for all the individuals, so it seems that there is certain ROH deficiency in coding regions. The situation was most striking for MHC-I-α, as the proportion of exons in ROH was much lower than expected (0.12 versus 0.41, i.e., 3.4x smaller on average), and highly significant for most individuals. There was also a significantly smaller proportion of OR exons in ROH for most individuals (0.28, i.e., 1.5x smaller on average; Table S12 and Figure 4B).

## Discussion

### Bloom filter assembly of a mammalian genome

Important progress has been made in genome sequencing technologies in recent years, leading to a decreased cost per base and a huge increase in the number of short sequences retrieved, allowing the *de novo* assembly of multiple species genomes with great coverage and quality (Goodwin et al. 2016). Nevertheless, the enormous quantity of data generated by these platforms has created new computational problems in terms of assembling large genomes, as this requires vast computational resources, especially memory (Sohn and Nam 2018). Of the algorithms that reduce overall memory requirements, Minia (Chikhi and Rizk 2013) and ABySS 2.0 (Jackman et al. 2017) use a Bloom filter to represent the de Bruijn graph, making it possible to assemble large genomes on low-memory computers. However, this Bloom filter-based approach has only been used in a few cases so far (Arnason et al. 2018; Renaut et al. 2018), probably because the method depends on a number of parameters that are not yet well understood and need to be tested. Here we show that the Bloom filter available in ABySS can be used to assemble the genome of the Pyrenean desman and produce a high-quality draft genome, with a scaffold N50 of 8.5 Mb and 96.3% of the BUSCO core genes. The final assembly was carried out in just 10 hours, using a computer with 128 GB of RAM memory and 16 processors. This reasonable timeframe and the possibility of running the program on a local computer allowed us to test many different settings, not only to properly tune the Bloom filter parameters but also to apply other conditions to obtain the best possible assembly. Part of the efficiency when assembling the Pyrenean desman genome could be related to the extremely low heterozygosity of the individual selected for the *de novo* assembly, which is one of the most important factors into obtaining a good assembly (Bradnam et al. 2013). However, the genome of the dromedary, with 710 SNPs/Mb (Fitak et al. 2016), was also adequately assembled, indicating that this method can be a good alternative for assembling large genomes, such as those of mammals, at very high coverage.

### Population demographic history of a species with low dispersal capacity

One of the most important life-history characteristics of the Pyrenean desman to help us understand the peculiar genomic features revealed in this work is its low dispersal capacity. The desman is morphologically well adapted to the aquatic medium, but its terrestrial locomotion is slow and labored (Palmeirim and Hoffmann 1983), meaning that, with a few exceptions, its dispersal occurs via the river network. Among the most important consequences of this low dispersal potential was the generation of a strong genetic structure during glacial periods, probably due to the complete isolation of glacial refugia over long periods, giving rise to five highly differentiated populations and strict contact zones with very low rates of mixing between adjacent populations (Igea et al. 2013; Querejeta et al. 2016; Escoda et al. 2017). During the period covered by the PSMC plot, there were important fluctuations in the size of these Pyrenean desman populations (Figure 2). Within a general trend of population decrease, two size peaks are apparent. Interestingly, they seem to coincide with the beginning of the two main interglacial periods of the last ∼300 thousand years. An expansion of the fluvial network during the deglaciations could have increased the favorable habitat for the Pyrenean desman and, consequently, its population. In addition, substantial differences between the demographic trajectories of the specimens sequenced were observed, much greater than the differences that are typically found between individuals of the same population (Nadachowska-Brzyska et al. 2016). These contrasting demographic histories are consistent with the different conditions likely to have been experienced by the Pyrenean desman populations during the glacial periods, and support the treatment of these populations as distinct evolutionarily significant units (Igea et al. 2013; Querejeta et al. 2016), which should be managed independently in conservation programs (Funk et al. 2012; Coates et al. 2018).

Range expansions and recurring bottlenecks in evolutionary history or recent past of a population can lead to a significant reduction in its genetic diversity (Hewitt 2000; Excoffier et al. 2009) and, consequently, the individual heterozygosity rate. The sequencing of species of conservation concern has led to comparisons being made between the heterozygosity rates of different species (Figure S6). Very rarely does a mammal have less than 100 SNPs/Mb; prior to this study, the lowest heterozygosity rate had been found in a Channel Island fox (*Urocyon littoralis*), with 14 SNPs/Mb, on the island of San Nicolas (Robinson et al. 2016). The Pyrenean desmans sequenced in this work span a wide range of heterozygosity rates, in line with the highly different evolutionary histories of the desman populations, with values running from 463 SNPs/Mb in the specimen from the West of the Iberian Peninsula to just 12 in the desman from the eastern Pyrenees. The latter is now the lowest heterozygosity rate recorded in any mammal, suggesting that the number of founding members of the population, situated at the eastern edge of the species range, could be as low as the number of foxes that colonized the small oceanic island of San Nicolas. The desmans from the western Pyrenees and the Central System are also positioned towards the bottom of the heterozygosity rate list (Figure S6), highlighting the ecological and evolutionary interest of these populations.

### Lessons from the genome of a species with extraordinary inbreeding levels

The reduced overland dispersal capacity of the Pyrenean desman has had profound effects on this species, not only during its recent history, but also in the present. Due to the abundance of artificial and ecological barriers in many of the rivers inhabited by this species, connectivity through the river network is currently greatly diminished. Large hydroelectric dams and water reservoirs very effectively block the movement of the desman. Additionally, the concatenation of smaller artificial barriers as well as ecological barriers resulting from contamination and predation by invasive species in the lower parts of rivers, has confined many desman populations to the river headwaters over the past few generations (Quaglietta et al. 2018). The consequence of this isolation is that desmans can only breed with other individuals of the same river, which are usually closely related as determined through relatedness networks (Escoda et al. 2017; Escoda et al. 2019). This, in turn, leads most desmans to have high inbreeding levels (Escoda et al. 2017). Considering that the inbreeding coefficient for the offspring of two first-degree relatives is 0.25 (Weir et al. 2006), values higher than this can only be achieved through continuous mating between closely related individuals for several generations. Five of the desmans sequenced in this study presented inbreeding coefficients greater than 0.25 and three of them even greater than 0.4 (Table S10; proportion of homozygous 100-kb windows). One of these is the desman from the Central System, which belongs to a highly isolated population recently discovered. Although the isolation of this population is mainly a result of ecological factors, the inbreeding coefficient of 0.49 found in the sequenced specimen is similar to those of desmans living in the headwaters above a large dam (Escoda et al. 2017). High inbreeding coefficients are also found in the two desmans from the western and eastern Pyrenees, with values of 0.47 and 0.69, respectively. However, the strong bottleneck experienced by these Pyrenean populations during the postglacial recolonization (Igea et al. 2013; Querejeta et al. 2016; Gillet et al. 2017) may also have affected the ROH. In fact, part of the ROH, the shortest runs, may be due to these more ancient phenomena and not only to recent inbreeding (Ceballos et al. 2018), thus explaining the extremely large proportion of ROH found in the Pyrenees.

The genomic sequences of individuals of an endangered species like the Pyrenean desman can also help determine the functional genomic features of particular specimens, to gain a better understanding of their genomic health (Steiner et al. 2013; Díez-del-Molino et al. 2018). In this work, we have characterized two groups of proteins from multigenic families in which high levels of diversity are essential, both at the inter- and intra-locus levels: the class I major histocompatibility complex (Hughes and Yeager 1998; Radwan et al. 2020) and the olfactory receptors (Alonso et al. 2008; Hughes et al. 2018). The analysis of the genetic diversity in these revealed interesting differences between the individuals sequenced. This was particularly true of the MHC-I-α genes, which must maintain high levels of genetic diversity to cope with external pathogens (Bateson et al. 2016; Marmesat et al. 2017). The desman from the West of the Iberian Peninsula and, to a certain extent, the two specimens from the Iberian System, maintain levels of heterozygosity in the MHC-I-α genes that are much higher than in other parts of the genome. However, in the three desmans from the Pyrenees and Central System these levels of intra-locus diversity are extremely low despite certain excess found, so the diversity of their immune system proteins can only be provided by genes from different loci. In principle, balancing selection could be acting in some populations to compensate for the sharp decrease occurred in heterozygosity throughout the genome due to the bottlenecks (Aguilar et al. 2004). However, we found that MHC-I-α genes tend to be absent from ROH regions, so this mechanism could also be important for maintaining genetic diversity where it is most necessary in highly inbred populations. A similar ROH deficiency in the MHC regions was found in the genome of cattle breeds (Zhang et al. 2015). The desman from the western Pyrenees is particularly interesting with regard to the MHC-I-α genes, as it has neither a high heterozygosity excess nor statistically significant reduction of these genes from ROH (Tables S11 and S12), suggesting that only inter-locus variability is maintaining its immune function. The OR genes also present a consistent heterozygosity excess in all desmans. The numbers of these genes are also reduced in ROH, particularly in the desman from the West of the Iberian Peninsula, in which almost no OR gene is present in ROH.

Therefore, an evolutionary mechanism through which MHC-I-α and OR genes are negatively selected in ROH regions may be acting. Since both MHC-I-α and OR genes are clustered in the genome, just a few regions could be targeted by this type of selection: individuals without ROH in them would have higher fitness and chances of surviving. Furthermore, this mechanism could be modulated by the proportion of deleterious mutations in the genome since their presence in ROH could be lethal, so perhaps the deficiency of certain genes in ROH can only be appreciated in species with low mutational load. Still, despite these mechanisms acting under current conditions, individuals with low overall genetic variability, particularly in the MCH genes, could be more susceptible to novel infectious diseases. Undoubtedly, a population genomics analysis with more individuals per population is necessary to thoroughly understand how different inbred specimens and populations cope with the need to maintain certain levels of genetic diversity in these important genes. This knowledge, together with the development of genomic health indicators based on the genetic diversity of key genes, could lead to a considerable improvement in *ex situ* management of threatened species, as only individuals with adequate indicators should be selected for genetic rescue or captive breeding. Further advances in this direction may open an interesting avenue of research with important applications for the conservation and management of endangered species.

A fundamental question that remains to be answered regarding the Pyrenean desman is whether these populations can survive with extremely low genome-wide heterozygosity, high proportion of ROH, and precariously maintained functional genetic diversity. Despite the shrinking habitat and range of this species (Fernandes et al. 2008), desmans are currently surviving with these poor genomic health indicators in the small river stretches to which the populations have become constrained. There is apparently no signal of generalized reduced fitness that may point to inbreeding depression, and new juveniles are detected every year, although we still do not know if some of these populations or all will collapse in the future. The reason why they continue to survive today may lie in a possible low mutational load. The bottlenecks experienced by the Pyrenean desman during the glaciations, as well as other adverse climatic periods such as droughts, could have purged deleterious and lethal mutations from the genomic background of the species, meaning that homozygosis is not as problematic today in the desman as it is in other species that present higher long-term genetic diversity but also more lethal equivalents (Keller and Waller 2002; Leberg and Firmin 2008). Even so, such low-diversity and highly inbred desman populations could be extremely vulnerable to the effects of pandemics caused by new pathogens, which may affect all individuals of the population equally (De Castro and Bolker 2004; Pedersen et al. 2007). Consequently, careful protection and monitoring of these populations is necessary. If population reinforcement becomes necessary, it should involve specimens from the same evolutionary unit and be planned with great caution because these genetically low-diversity populations might be particularly difficult to rescue, as there are high chances of introducing elevated levels of recessive mutations from large populations (Kyriazis et al. 2019). For this reason, any conservation strategies should preferentially promote natural connectivity between nearby river populations or, where this is not feasible, proceed with reciprocal translocations between recently disconnected populations. Genomics can help to not only determine which specimens may be more or less appropriate for genetic rescue or captive breeding according to specific genomic health indicators of each individual (Supple and Shapiro 2018; Humble et al. 2020; Wright et al. 2020), but also to monitor future individuals sampled after the conservation actions to confirm if the measures employed are helping to improve the impoverished genomic health of the Pyrenean desman.

## Materials and Methods

### Genome sequencing and assembly

We selected tissue samples from six Pyrenean desmans, two of which (IBE-C3734 and IBE-C3773) had been utilized in a previous ddRAD study (Escoda et al. 2017), and extracted DNA using the methods described in that report (Table S1). All the samples used in this study were minimally invasive samples obtained as part of works with the species promoted by environmental authorities or came from animals found dead during these surveys. The DNA quality of the samples selected for genome sequencing was controlled by checking for the absence of smearing in a gel electrophoresis. The Pyrenean desman from the West of the Iberian Peninsula (IBE-C6507) was sequenced using ddRAD (Peterson et al. 2012) and analyzed together with previously sequenced samples from the same population (Escoda et al. 2019) in order to estimate its inbreeding coefficient using a maximum-likelihood method based on the population frequencies (Wang 2011). Its heterozygosity rate was calculated from the ddRAD sequences, which represent approximately 0.1 % of the genome.

A single male specimen (IBE-C5619) was used for sequencing the reference genome. Its genomic DNA was shotgun-sequenced using three Illumina TruSeq DNA PCR-free libraries, two with an insert size of 350 bp and one of 550 bp, and two mate-pair libraries with insert sizes of 5 kb and 9 kb, respectively. For the five additional resequenced genomes, a TruSeq DNA PCR-free library with an insert size of 350 bp was constructed for each individual. All the libraries were prepared by Macrogen Inc. (South Korea) and sequenced in either Illumina HiSeq X Ten or Novaseq 6000 instruments.

We separated the different fractions of the mate-pair reads with NxTrim v0.4.3 (O’Connell et al. 2015) and for the assembly we used only the fraction with mate-pair orientation and complete reads, i.e., with no adapter sequence (called unknown in NxTrim), which produced the best results in initial assemblies. Then, we used fastp v0.19.5 (Chen et al. 2018) with all the libraries to remove adapters and low complexity reads (to eliminate sequencing artifacts), as well as reads with a quality score of lower than 20 or a length of less than 150 bp. Using the same tool, the reads were base-corrected.

Using the filtered reads, we assembled the *de novo* genome using ABySS v2.1.5 (Jackman et al. 2017). We applied the Bloom filter option for contig formation and tested several parameters of both contigs and scaffolds formation stages: k-mer size (k), minimum k-mer count threshold for Bloom filter assembly (kc), minimum number of pairs required for building contigs (n), and minimum number of pairs required for building scaffolds (N). All the assemblies were carried out with the following parameters in common, as preliminary analyses showed them to be the best: Bloom filter size (B) of 80G, number of Bloom filter hash functions (H) of 4, and minimum untig size required for building contigs (s) of 1000.

We used QUAST v5.0.2 (Gurevich et al. 2013) to compute the summary statistics and BUSCO (Benchmarking Universal Single-Copy Orthologs) v3.0.2 (Simão et al. 2015) to assess the genome completeness of the different assemblies. The best assembly parameters were chosen in order to maximize the N50 of the assembly and the number of core genes found with BUSCO, as well as to minimize the number of scaffolds and gaps (N’s). The GC content was calculated using BEDTools (Quinlan and Hall 2010) in 100-kb windows.

In order to assess the assembly performed with the Bloom filter option, we downloaded the genome and raw sequences of a dromedary already assembled using ABySS (Fitak et al. 2016), and compared this to the assembly using the Bloom filter option. To do this, we filtered the raw reads of both the paired-end and mate-pair libraries with fastp v0.19.5 (Chen et al. 2018), as before, and assembled them using the Bloom filter option in ABySS with a k-mer size of 64 (different from the desman assembly due to the shorter reads of the dromedary); for the remaining parameters we applied the best options found for the desman assembly. To compare the two assemblies, we masked the repetitive regions of both, as detailed below, aligned the scaffolds longer than 10,000 bp using MUMmer v3.23 (Kurtz et al. 2004) with the nucmer option and parameters “-maxmatch -l 100 -c 100”, and plotted the alignment with mummerplot after filtering for the alignments that represented the best one-to-one mapping of both genomes.

### Gene prediction

We identified the repetitive regions in the genome assembly with RepeatMasker v4.0.7 (http://www.repeatmasker.org) using the Dfam Consensus release 20170127 and RepBase release 20170127 databases (Jurka et al. 2005). Complex repeats were hard-masked whereas simple repeats were soft-masked so that they could be used in some further steps.

For gene detection, we used two iterative rounds of the MAKER2 v2.31.10 pipeline (Holt and Yandell 2011) with the masked genome sequence (with contigs >= 1,000 bp). In the first round, genes were predicted using two methods: directly from protein homology (option protein2genome=1) using exonerate (Slater and Birney 2005); and also with AUGUSTUS v3.3.2 (Stanke et al. 2006), previously trained with a small fraction of the genome. For the protein homology prediction, we used the proteomes from the only four species of the Eulipotyphla order to which the Pyrenean desman belongs: *Condylura cristata, Sorex araneus*, and *Erinaceus europaeus*, all of them unpublished genomes from the Broad Institute available at GenBank (Clark et al. 2016), and *Solenodon paradoxus* (Casewell et al. 2019). In addition, we included the human proteome available at GenBank, as its completeness allowed us to detect additional genes. These proteomes were also used as protein evidence, as well as to refine the gene models using exonerate. In the second round, genes were predicted using AUGUSTUS, as before, and also with SNAP version 2013-02-16 (Korf 2004), the latter trained with predictions of the first MAKER2 round. For the two rounds, an expected maximum intron size of 15,000 bp was used and scaffolds were divided into chunks of 400,000 bp. Gene annotation of the generated GFF and FASTA files was based on a BLAST search (Altschul et al. 1997) against the mammal section of the UniProt/Swiss-Prot database (UniProt Consortium 2019). GenomeTools v1.5.10 (Gremme et al. 2013) was used to compute statistics on the predicted genes.

MHC-I-α genes were retrieved according to the corresponding annotations (”class I histocompatibility antigen” and “alpha chain”) from the GFF and FASTA files. Out of 45 genes found, only 26 with between 5 and 8 exons were retained. For the heterozygous sequences, we selected the assembled sequence. Alignments at the amino acid level of these sequences together with those from other mammals (Abduriyim et al. 2019), including *Condylura cristata* and a selection of human genes, were generated with MAFFT v7.464 (Katoh and Standley 2013) and processed with Gblocks v0.91 (Castresana 2000) to remove poorly aligned positions using low stringency conditions (minimum length of a block of 5 and allowing positions with gaps in half the number of sequences). Then, a maximum-likelihood phylogenetic tree was reconstructed with RAxML v8.2.12, using a JTT model of amino acid substitution and a gamma distribution of evolutionary rates (Stamatakis 2014).

To identify olfactory receptor (OR) genes, we used 659 putative OR genes obtained from the first round of MAKER2 annotated as “olfactory receptor” in the GFF file, as this round contained a much larger number of putative OR genes obtained by protein homology. We classified this set of putative OR genes into functional genes and non-functional pseudogenes with the Olfactory Receptor family Assigner (ORA) BioPerl module (Hayden et al. 2010). In addition, four genes with large numbers of masked sequences and a gene that was very divergent in initial trees were eliminated. The alignment and phylogenetic tree of the final 529 genes together with those from *Condylura cristata* and human were generated as described above.

### Read mapping and variant calling

Cleaned reads of each individual were mapped to the *de novo* reference genome using BWA v0.7.17 (Li and Durbin 2009). Subsequently, SAMtools v1.9 (Li et al. 2009) was used to produce BAM alignments of scaffold length greater than 1,000 bp in which duplicated reads were removed and only unambiguously mapped and properly paired reads with a minimum mapping quality of 20 were kept. Variant calling was carried out with BCFtools v1.9 (Li 2011) from the BAM alignments in scaffolds with a minimum length of 40,000 bp. This threshold was selected because we observed that, according to the result of QualiMap v2.2.2 (Okonechnikov et al. 2016), the mapping quality and insert size graphs greatly improved in scaffolds longer than this length, including their two ends (where mapping quality was reduced in smaller scaffolds), something essential for proper genetic diversity estimates. Additional filtering parameters for obtaining the final VCF files of the genotypes included a minimum variant quality of 30, a maximum depth of coverage of twice the average of the genome-wide coverage of each individual, and a minimum depth of coverage of either 5 or 12, depending on the analysis.

### Genomic heterozygosity

Genome-wide heterozygosity was based on the genotypes of each individual obtained after discarding SNPs due to differences with the reference genome, therefore keeping only heterozygous positions. The number of heterozygous positions was obtained with VCFtools v0.1.16 (Danecek et al. 2011) from the VCF files of each individual based on a minimum depth of coverage of either 5 or 12. This was then divided by the number of positions that passed equivalent filters from the corresponding BAM files, as calculated with SAMtools, to estimate the heterozygosity rate.

To detect sexual scaffolds, we computed the heterozygosity of scaffolds longer than 40,000 bp, as before, using the genotype calls with a minimum depth of coverage of 12. We based the classification of the chromosomes on the fact that the ratio of coverage between the female and any of the males presented three clearly delimited groups: autosomes (ratio ∼ 1), X chromosomes (∼ 2), and Y chromosomes (∼ 0). We considered those with a ratio of coverage of between 0 and 0.04 to be Y-chromosome scaffolds, and those with a ratio between 1.5 and 2.5 to be X-chromosome scaffolds. After excluding Y- and X-chromosome scaffolds, 583 putative autosomal scaffolds longer than 40,000 bp remained. These autosomal scaffolds were the basis of further analyses, including the final computation of genome-wide heterozygosity described above.

The number of heterozygous positions in exons of different genes was calculated with BEDtools (Quinlan and Hall 2010) by crossing the VCF files of heterozygosity for each individual (with a minimum depth of coverage of 5) with the BED files containing the exon positions of the desired genes, all in the 583 autosomal scaffolds longer than 40,000 bp. The resulting number of heterozygous positions was divided by the total number of exon positions calculated from the corresponding exon BED files to estimate the heterozygosity rate in each gene class.

### Runs of homozygosity (ROH)

We identified ROH using four different approaches. Firstly, we used BCFtools/RoH (Narasimhan et al. 2016), which applies a hidden Markov model, applying default parameters except that the window size was 100 kb. Secondly, we used PLINK v1.90p (Purcell et al. 2007) to detect ROH segments larger than 100 kb with default options, in which a scanning window of 50 SNPs can contain, at most, 1 heterozygous position. These two methods are based on the variable sites present across the six sequenced individuals and we used the VCF file from all the individuals obtained with a minimum depth of coverage of 5. For the final calculation of the ROH proportion of the genome, we only considered the fraction of the genome in scaffolds longer than 100 kb.

We also used two additional methods that are based on each individual genome for identifying ROH. We first used ROHan (Renaud et al. 2019), which simultaneously estimates heterozygosity and identifies ROH regions from the BAM file of each individual genome, using 100-kb windows and allowing 0.5 heterozygous position per 100 kb. Although the default is a maximum of 1 heterozygous position per 100 kb, this rate (10 SNPs/Mb) is too high for desman populations where this value is close to the genome average. Finally, we calculated the heterozygosity rate of the autosomal scaffolds with VCFtools v0.1.16 (Danecek et al. 2011), in 100-kb non-overlapping windows, using the VCF files of heterozygosity for each individual with a minimum depth of coverage of 5. Then, we calculated the proportion of 100-kb windows that contained 0 heterozygous positions, which were considered ROH windows, with respect to all the 100-kb windows (not including partial windows).

The proportion of exons of MHC-I-α and OR in ROH was calculated with BEDtools (Quinlan and Hall 2010) by means of crossing the BED files containing the exon positions of the desired genes with the BED files of ROH and non-ROH 100-kb windows of each individual. The proportion was calculated as the number of exons in ROH windows divided by the total number of exons in 100-kb windows. To calculate the p-value, the ROH and non-ROH 100-kb windows of each individual were randomized 1000 times while maintaining the proportion of windows with 0 heterozygous positions corresponding to each individual.

### Genetic structure and demographic history

To assemble the mitochondrial genomes, we mapped the raw reads of each individual to a published complete mitochondrial genome (Cabria et al. 2006) using BWA v0.7.17 (Li and Durbin 2009) and called variants using BCFtools v1.9 (Li 2011). We obtained each mitochondrial genome by applying the variants of each individual to the reference mitogenome with the BCFtools consensus tool. All the mitogenomes were aligned using MAFFT (Katoh and Standley 2013) and a maximum-likelihood phylogenetic tree was reconstructed with RAxML using a GTR model of nucleotide substitution and a gamma distribution of evolutionary rates (Stamatakis 2014).

PCA was performed with the KING toolset (Manichaikul et al. 2010) using the genotypes obtained with a minimum depth of coverage of 5.

For the PSMC analysis (Li and Durbin 2011), we first used the genotypes with a minimum depth of coverage of 5 to generate a consensus FASTA sequence of the autosomal genome scaffolds. We then performed the PSMC analyses using the following parameters, as suggested in the program manual (https://github.com/lh3/psmc): maximum number of iterations (N) of 25, maximum coalescent time (t) of 15, initial theta/rho ratio (r) of 5, and parameter pattern (p) of “4+25*2+4+6”. The program infers the recombination rate and the population size parameters. Similar results were found when using the alternative parameters: N=25, t=5, r=1, and p=”4+30*2+4+6+10” (Nadachowska-Brzyska et al. 2016). We assessed the variance of the analyses using 100 bootstrap replicates of each individual. The final estimates of population size and time were scaled with a mutation rate of 5 × 10^−9^ mutations/site/generation and a generation time of 2 years. The Pyrenean desman can live up to 4 years, and occasionally as long as 6 (Gonzalez-Esteban et al. 2002), while reconstructed pedigrees (Escoda et al. 2019) suggest that 2 years approximates well to the average intergeneration interval. The mutation rate per generation of 5 × 10^−9^ was selected as this is similar to other species with short generation times (Uchimura et al. 2015; Smeds et al. 2016), as expected (Piganeau and Eyre-Walker 2009); the resulting per year mutation rate for the desman (2.5 × 10^−9^ mutations/site/year) was similar to the mammalian average of 2.2 × 10^−9^ mutations/site/year (Kumar and Subramanian 2002).

## Data Availability

Upon acceptance for publication in a peer-reviewed journal, data will be deposited in a digital repository.

## Supplementary Material

Additional data and figures are available in Supplementary Material.

## Acknowledgements

We thank Ángel Fernández-González, Jorge González-Esteban, Pere Aymerich, Núria Valls-Granero, Oriol Comas-Angelet and people from Biosfera Consultoría Medioambiental S.L. for providing samples of Pyrenean desmans as well as for sharing information on the biology, ecology and conservation issues of this species, and Julio Rozas for critically reading the manuscript and useful suggestions. We also thank Junta de Castilla y León, Generalitat de Catalunya, Gobierno de Navarra, and Gobierno de La Rioja for permits to use samples in our studies and particularly David Cubero, Gabriel de Pedro, and Sisco Mañas for their help. This work was supported by research Project CGL2017-84799-P of the “Plan Nacional I+D+i del Ministerio de Ciencia e Innovación” (Spain), cofinanced with FEDER funds, to J.C.

## References

Abascal F, Corvelo A, Cruz F, Villanueva-Cañas JL, Vlasova A, Marcet-Houben M, Martínez-Cruz B, Cheng JY, Prieto P, Quesada V, et al. 2016. Extreme genomic erosion after recurrent demographic bottlenecks in the highly endangered Iberian lynx. Genome Biol. 17:251.

Abduriyim S, Zou DH, Zhao H. 2019. Origin and evolution of the major histocompatibility complex class I region in eutherian mammals. Evol. Ecol. 9:7861–7874.

Aguilar A, Roemer G, Debenham S, Binns M, Garcelon D, Wayne RK. 2004. High MHC diversity maintained by balancing selection in an otherwise genetically monomorphic mammal. Proc. Natl Acad. Sci. USA 101:3490–3494.

Alonso S, López S, Izagirre N, la Rúa de C. 2008. Overdominance in the human genome and olfactory receptor activity. Mol. Biol. Evol. 25:997–1001.

Altschul SF, Madden TL, Schäffer AA, Zhang J, Zhang Z, Miller W, Lipman DJ. 1997. Gapped BLAST and PSI-BLAST: a new generation of protein database search programs. Nucleic Acids Res. 25:3389–3402.

Arnason U, Lammers F, Kumar V, Nilsson MA, Janke A. 2018. Whole-genome sequencing of the blue whale and other rorquals finds signatures for introgressive gene flow. Science Advances 4:eaap9873.

Bateson ZW, Hammerly SC, Johnson JA, Morrow ME, Whittingham LA, Dunn PO. 2016. Specific alleles at immune genes, rather than genome-wide heterozygosity, are related to immunity and survival in the critically endangered Attwater’s prairie-chicken. Mol. Ecol. 25:4730–4744.

Benazzo A, Trucchi E, Cahill JA, Maisano Delser P, Mona S, Fumagalli M, Bunnefeld L, Cornetti L, Ghirotto S, Girardi M, et al. 2017. Survival and divergence in a small group: The extraordinary genomic history of the endangered Apennine brown bear stragglers. Proc. Natl Acad. Sci. USA 114:E9589–E9597.

Bradnam KR, Fass JN, Alexandrov A, Baranay P, Bechner M, Birol I, Boisvert S, Chapman JA, Chapuis G, Chikhi R, et al. 2013. Assemblathon 2: evaluating *de novo* methods of genome assembly in three vertebrate species. GigaScience 2:10.

Cabria MT, Rubines J, Gómez-Moliner BJ, Zardoya R. 2006. On the phylogenetic position of a rare Iberian endemic mammal, the Pyrenean desman (*Galemys pyrenaicus*). Gene 375:1–13.

Campbell MS, Holt C, Moore B, Yandell M. 2014. Genome Annotation and Curation Using MAKER and MAKER-P. Curr. Protoc. Bioinformatics 48:4.11.1–4.11.39.

Casewell NR, Petras D, Card DC, Suranse V, Mychajliw AM, Richards D, Koludarov I, Albulescu L-O, Slagboom J, Hempel B-F, et al. 2019. Solenodon genome reveals convergent evolution of venom in eulipotyphlan mammals. Proc. Natl Acad. Sci. USA 116:25745–25755.

Castresana J. 2000. Selection of conserved blocks from multiple alignments for their use in phylogenetic analysis. Mol. Biol. Evol. 17:540–552.

Ceballos FC, Joshi PK, Clark DW, Ramsay M, Wilson JF. 2018. Runs of homozygosity: windows into population history and trait architecture. Nat. Rev. Genet. 19:220–234.

Chen S, Zhou Y, Chen Y, Gu J. 2018. fastp: an ultra-fast all-in-one FASTQ preprocessor. Bioinformatics 34:i884–i890.

Chikhi R, Rizk G. 2013. Space-efficient and exact de Bruijn graph representation based on a Bloom filter. Algorithms Mol Biol 8:22.

Clark K, Karsch-Mizrachi I, Lipman DJ, Ostell J, Sayers EW. 2016. GenBank. Nucleic Acids Res. 44:D67–D72.

Clark PU, Dyke AS, Shakun JD, Carlson AE, Clark J, Wohlfarth B, Mitrovica JX, Hostetler SW, McCabe AM. 2009. The Last Glacial Maximum. Science 325:710–714.

Coates DJ, Byrne M, Moritz CC. 2018. Genetic Diversity and Conservation Units: Dealing With the Species-Population Continuum in the Age of Genomics. Front. Ecol. Evol. 6:4045.

Dahl-Jensen D, Albert MR, Aldahan A, Azuma N, Balslev-Clausen D, Baumgartner M, Berggren AM, Bigler M, Binder T, Blunier T, et al. 2013. Eemian interglacial reconstructed from a Greenland folded ice core. Nature 493:489–494.

Danecek P, Auton A, Abecasis G, Albers CA, Banks E, DePristo MA, Handsaker RE, Lunter G, Marth GT, Sherry ST, et al. 2011. The variant call format and VCFtools. Bioinformatics 27:2156–2158.

De Castro F, Bolker B. 2004. Mechanisms of disease-induced extinction. Ecol. Lett. 8:117–126.

Díez-del-Molino D, Sánchez Barreiro F, Barnes I, Gilbert MTP, Dalén L. 2018. Quantifying Temporal Genomic Erosion in Endangered Species. Trends Ecol. Evol. 33:176–185.

Ekblom R, Brechlin B, Persson J, Smeds L, Johansson M, Magnusson J, Flagstad Ø, Ellegren H. 2018. Genome sequencing and conservation genomics in the Scandinavian wolverine population. Conserv Biol 32:1301–1312.

Escoda L, Fernández-González A, Castresana J. 2019. Quantitative analysis of connectivity in populations of a semi-aquatic mammal using kinship categories and network assortativity. Mol. Ecol. Resour. 19:310–326.

Escoda L, Gonzalez-Esteban J, Gómez A, Castresana J. 2017. Using relatedness networks to infer contemporary dispersal: Application to the endangered mammal *Galemys pyrenaicus*. Mol. Ecol. 26:3343–3357.

Excoffier L, Foll M, Petit R. 2009. Genetic Consequences of Range Expansions. Annu. Rev. Ecol. Evol. Syst. 40:481–501.

Fernandes M, Herrero J, Aulagnier S, Amori G. 2008. Galemys pyrenaicus. IUCN Red List of Threatened Species:e.T8826A12934876.

Fitak RR, Mohandesan E, Corander J, Burger PA. 2016. The *de novo* genome assembly and annotation of a female domestic dromedary of North African origin. Mol. Ecol. Resour. 16:314–324.

Funk WC, Mckay JK, Hohenlohe PA, Allendorf FW. 2012. Harnessing genomics for delineating conservation units. Trends Ecol. Evol. 27:489–496.

Gillet F, Cabria Garrido MT, Blanc F, Fournier-Chambrillon C, N moz M, Sourp E, Vial-Novella C, Zardoya R, Aulagnier S, Michaux JR. 2017. Evidence of fine-scale genetic structure for the endangered Pyrenean desman (*Galemys pyrenaicus*) in the French Pyrenees. J. Mammal. 98:523–532.

Gonzalez-Esteban J, Villate I, Castién E, Rey I, Gosálbez J. 2002. Age determination of *Galemys pyrenaicus*. Acta Theriol. 47:107–112.

Goodwin S, McPherson JD, McCombie WR. 2016. Coming of age: ten years of next-generation sequencing technologies. Nat. Rev. Genet. 17:333–351.

Gremme G, Steinbiss S, Kurtz S. 2013. GenomeTools: a comprehensive software library for efficient processing of structured genome annotations. IEEE/ACM Transactions on Computational Biology and Bioinformatics 10:645–656.

Gurevich A, Saveliev V, Vyahhi N, Tesler G. 2013. QUAST: quality assessment tool for genome assemblies. Bioinformatics 29:1072–1075.

Hayden S, Bekaert M, Crider TA, Mariani S, Murphy WJ, Teeling EC. 2010. Ecological adaptation determines functional mammalian olfactory subgenomes. Genome Res. 20:1–9.

Hewitt GM. 2000. The genetic legacy of the Quaternary ice ages. Nature 405:907–913.

Holt C, Yandell M. 2011. MAKER2: an annotation pipeline and genome-database management tool for second-generation genome projects. BMC Bioinformatics 12:491.

Hughes AL, Yeager M. 1998. Natural selection at major histocompatibility complex loci of vertebrates. Annu Rev Genet 32:415–435.

Hughes GM, Boston ESM, Finarelli JA, Murphy WJ, Higgins DG, Teeling EC. 2018. The Birth and Death of Olfactory Receptor Gene Families in Mammalian Niche Adaptation. Mol. Biol. Evol. 35:1390–1406.

Humble E, Dobrynin P, Senn H, Chuven J, Scott AF, Mohr DW, Dudchenko O, Omer AD, Colaric Z, Lieberman Aiden E, et al. 2020. Chromosomal-level genome assembly of the scimitar-horned oryx: Insights into diversity and demography of a species extinct in the wild. Mol. Ecol. Resour. doi: 10.1111/1755-0998.13181.

Igea J, Aymerich P, Fernández-González A, Gonzalez-Esteban J, Gómez A, Alonso R, Gosálbez J, Castresana J. 2013. Phylogeography and postglacial expansion of the endangered semi-aquatic mammal *Galemys pyrenaicus*. BMC Evol. Biol. 13:115.

Jackman SD, Vandervalk BP, Mohamadi H, Chu J, Yeo S, Hammond SA, Jahesh G, Khan H, Coombe L, Warren RL, et al. 2017. ABySS 2.0: resource-efficient assembly of large genomes using a Bloom filter. Genome Res. 27:768–777.

Jurka J, Kapitonov VV, Pavlicek A, Klonowski P, Kohany O, Walichiewicz J. 2005. Repbase Update, a database of eukaryotic repetitive elements. Cytogenet. Genome Res. 110:462–467.

Kardos M, Taylor HR, Ellegren H, Luikart G, Allendorf FW. 2016. Genomics advances the study of inbreeding depression in the wild. Evol. Appl. 9:1205–1218.

Katoh K, Standley DM. 2013. MAFFT multiple sequence alignment software version 7: improvements in performance and usability. Mol. Biol. Evol. 30:772–780.

Keller LF, Waller DM. 2002. Inbreeding effects in wild populations. Trends Ecol. Evol. 17:230–241.

Korf I. 2004. Gene finding in novel genomes. BMC Bioinformatics 5:59.

Kryštufek B, Motokawa M. 2018. Species accounts of Talpidae. In: Mittermeier RA, Wilson DE, editors. Handbook of the Mammals of the World. Volume 8. Insectivores, Sloths and Colugos. Barcelona (Spain): Lynx Edicions.

Kumar S, Subramanian S. 2002. Mutation rates in mammalian genomes. Proc. Natl Acad. Sci. USA 99:803–808.

Kurtz S, Phillippy A, Delcher AL, Smoot M, Shumway M, Antonescu C, Salzberg SL. 2004. Versatile and open software for comparing large genomes. Genome Biol. 5:R12.

Kyriazis CC, Wayne RK, Lohmueller KE. 2019. High genetic diversity can contribute to extinction in small populations. bioRxiv 10.1101/678524.

Labeit S, Kolmerer B. 1995. Titins: giant proteins in charge of muscle ultrastructure and elasticity. Science 270:293–296.

Leberg PL, Firmin BD. 2008. Role of inbreeding depression and purging in captive breeding and restoration programmes. Mol. Ecol. 17:334–343.

Li H, Durbin R. 2009. Fast and accurate short read alignment with Burrows-Wheeler transform. Bioinformatics 25:1754–1760.

Li H, Durbin R. 2011. Inference of human population history from individual whole-genome sequences. Nature 475:493–496.

Li H, Handsaker B, Wysoker A, Fennell T, Ruan J, Homer N, Marth G, Abecasis G, Durbin R, 1000 Genome Project Data Processing Subgroup. 2009. The Sequence Alignment/Map format and SAMtools. Bioinformatics 25:2078–2079.

Li H. 2011. A statistical framework for SNP calling, mutation discovery, association mapping and population genetical parameter estimation from sequencing data. Bioinformatics 27:2987–2993.

Locke DP, Hillier LW, Warren WC, Worley KC, Nazareth LV, Muzny DM, Yang S-P, Wang Z, Chinwalla AT, Minx P, et al. 2011. Comparative and demographic analysis of orang-utan genomes. Nature 469:529–533.

Manichaikul A, Mychaleckyj JC, Rich SS, Daly K, Sale M, Chen W-M. 2010. Robust relationship inference in genome-wide association studies. Bioinformatics 26:2867–2873.

Marmesat E, Schmidt K, Saveljev AP, Seryodkin IV, Godoy JA. 2017. Retention of functional variation despite extreme genomic erosion: MHC allelic repertoires in the *Lynx* genus. BMC Evol. Biol. 17:158.

Nadachowska-Brzyska K, Burri R, Smeds L, Ellegren H. 2016. PSMC analysis of effective population sizes in molecular ecology and its application to black-and-white *Ficedula* flycatchers. Mol. Ecol. 25:1058–1072.

Narasimhan V, Danecek P, Scally A, Xue Y, Tyler-Smith C, Durbin R. 2016. BCFtools/RoH: a hidden Markov model approach for detecting autozygosity from next-generation sequencing data. Bioinformatics 32:1749–1751.

O’Connell J, Schulz-Trieglaff O, Carlson E, Hims MM, Gormley NA, Cox AJ. 2015. NxTrim: optimized trimming of Illumina mate pair reads. Bioinformatics 31:2035–2037.

Okonechnikov K, Conesa A, García-Alcalde F. 2016. Qualimap 2: advanced multi-sample quality control for high-throughput sequencing data. Bioinformatics 32:292–294.

Palmeirim JM, Hoffmann RS. 1983. *Galemys pyrenaicus*. Mamm. Species 207:1–5.

Pedersen AB, Jones KE, Nunn CL, Altizer S. 2007. Infectious Diseases and Extinction Risk in Wild Mammals. Conserv Biol 21:1269–1279.

Peterson BK, Weber JN, Kay EH, Fisher HS, Hoekstra HE. 2012. Double digest RADseq: an inexpensive method for *de novo* SNP discovery and genotyping in model and non-model species. PLOS ONE 7:e37135.

Piganeau G, Eyre-Walker A. 2009. Evidence for variation in the effective population size of animal mitochondrial DNA. PLOS ONE 4:e4396.

Piovesan A, Caracausi M, Antonaros F, Pelleri MC, Vitale L. 2016. GeneBase 1.1: a tool to summarize data from NCBI gene datasets and its application to an update of human gene statistics. Database 2016:1–13.

Prado-Martinez J, Sudmant PH, Kidd JM, Li H, Kelley JL, Lorente-Galdos B, Veeramah KR, Woerner AE, O’Connor TD, Santpere G, et al. 2013. Great ape genetic diversity and population history. Nature 499:471–475.

Purcell S, Neale B, Todd-Brown K, Thomas L, Ferreira MAR, Bender D, Maller J, Sklar P, de Bakker PIW, Daly MJ, et al. 2007. PLINK: a tool set for whole-genome association and population-based linkage analyses. Am. J. Hum. Genet. 81:559–575.

Quaglietta L, Pauperio J, Martins FM, Alves PC, Beja P. 2018. Recent range contractions in the globally threatened Pyrenean desman highlight the importance of stream headwater refugia. Anim Conserv 21:515–525.

Querejeta M, Gonzalez-Esteban J, Gómez A, Fernández-González A, Aymerich P, Gosálbez J, Escoda L, Igea J, Castresana J. 2016. Genomic diversity and geographical structure of the Pyrenean desman. Conserv. Genet. 17:1333–1344.

Quinlan AR, Hall IM. 2010. BEDTools: a flexible suite of utilities for comparing genomic features. Bioinformatics 26:841–842.

Radwan J, Babik W, Kaufman J, Lenz TL, Winternitz J. 2020. Advances in the Evolutionary Understanding of MHC Polymorphism. Trends Genet. 36:298–311.

Renaud G, Hanghøj K, Korneliussen TS, Willerslev E, Orlando L. 2019. Joint Estimates of Heterozygosity and Runs of Homozygosity for Modern and Ancient Samples. Genetics 212:587–614.

Renaut S, Guerra D, Hoeh WR, Stewart DT, Bogan AE, Ghiselli F, Milani L, Passamonti M, Breton S. 2018. Genome Survey of the Freshwater Mussel *Venustaconcha ellipsiformis* (Bivalvia: Unionida) Using a Hybrid De Novo Assembly Approach. Genome Biology and Evolution 10:1637–1646.

Robinson JA, Ortega-Del Vecchyo D, Fan Z, Kim BY, Vonholdt BM, Marsden CD, Lohmueller KE, Wayne RK. 2016. Genomic Flatlining in the Endangered Island Fox. Curr. Biol. 26:1183–1189.

Saremi NF, Supple MA, Byrne A, Cahill JA, Coutinho LL, Dalén L, Figueiró HV, Johnson WE, Milne HJ, O’Brien SJ, et al. 2019. Puma genomes from North and South America provide insights into the genomic consequences of inbreeding. Nat Commun 10:4769.

Simão FA, Waterhouse RM, Ioannidis P, Kriventseva EV, Zdobnov EM. 2015. BUSCO: assessing genome assembly and annotation completeness with single-copy orthologs. Bioinformatics 31:3210–3212.

Slater GSC, Birney E. 2005. Automated generation of heuristics for biological sequence comparison. BMC Bioinformatics 6:31.

Smeds L, Qvarnström A, Ellegren H. 2016. Direct estimate of the rate of germline mutation in a bird. Genome Res. 26:1211–1218.

Sohn J-I, Nam J-W. 2018. The present and future of de novo whole-genome assembly. Brief. Bioinformatics 19:23–40.

Stamatakis A. 2014. RAxML version 8: a tool for phylogenetic analysis and post-analysis of large phylogenies. Bioinformatics 30:1312–1313.

Stanke M, Keller O, Gunduz I, Hayes A, Waack S, Morgenstern B. 2006. AUGUSTUS: ab initio prediction of alternative transcripts. Nucleic Acids Res. 34:W435–W439.

Steiner CC, Putnam AS, Hoeck PEA, Ryder OA. 2013. Conservation genomics of threatened animal species. Annu Rev Anim Biosci 1:261–281.

Supple MA, Shapiro B. 2018. Conservation of biodiversity in the genomics era. Genome Biol. 19:131.

Uchimura A, Higuchi M, Minakuchi Y, Ohno M, Toyoda A, Fujiyama A, Miura I, Wakana S, Nishino J, Yagi T. 2015. Germline mutation rates and the long-term phenotypic effects of mutation accumulation in wild-type laboratory mice and mutator mice. Genome Res. 25:1125–1134.

UniProt Consortium. 2019. UniProt: a worldwide hub of protein knowledge. Nucleic Acids Res. 47:D506–D515.

Vandiedonck C, Knight JC. 2009. The human Major Histocompatibility Complex as a paradigm in genomics research. Briefings in Functional Genomics and Proteomics 8:379–394.

Wang J. 2011. COANCESTRY: a program for simulating, estimating and analysing relatedness and inbreeding coefficients. Mol. Ecol. Resour. 11:141–145.

Weir BS, Anderson AD, Hepler AB. 2006. Genetic relatedness analysis: modern data and new challenges. Nat. Rev. Genet. 7:771–780.

Westbury MV, Hartmann S, Barlow A, Wiesel I, Leo V, Welch R, Parker DM, Sicks F, Ludwig A, Dalén L, et al. 2018. Extended and Continuous Decline in Effective Population Size Results in Low Genomic Diversity in the World’s Rarest Hyena Species, the Brown Hyena. Mol. Biol. Evol. 35:1225–1237.

Wright BR, Farquharson KA, McLennan EA, Belov K, Hogg CJ, Grueber CE. 2020. A demonstration of conservation genomics for threatened species management. Mol. Ecol. Resour. doi: 10.1111/1755-0998.13211.

Xue Y, Prado-Martinez J, Sudmant PH, Narasimhan V, Ayub Q, Szpak M, Frandsen P, Chen Y, Yngvadottir B, Cooper DN, et al. 2015. Mountain gorilla genomes reveal the impact of longterm population decline and inbreeding. Science 348:242–245.

Zhang Q, Guldbrandtsen B, Bosse M, Lund MS, Sahana G. 2015. Runs of homozygosity and distribution of functional variants in the cattle genome. BMC Genomics 16:542.

