## Supplementary material for "The impact of bottlenecks and inbreeding on the genome of the endangered Pyrenean desman": Tables and Figures

**Index of Supplementary Information**

**Table S1**. Specimens used in this study, sex, sampling year, locality, and geographical area.
**Table S2**. Summary statistics of the *de novo* genome sequencing data.
**Table S3**. Summary statistics of the genome sequencing data of the additional individuals.
**Table S4**. Summary statistics of the genome assemblies using different combinations of parameters in ABySS.
**Table S5**. Summary statistics of the genome assembly.
**Table S6**. Summary of the BUSCO analysis.
**Table S7**. Summary statistics of the genome assemblies of the dromedary using ABySS.
**Table S8**. Summary statistics of the repetitive elements.
**Table S9**. Autosomal genome-wide heterozygosity of the sequenced Pyrenean desmans.
**Table S10**. Runs of homozygosity (ROH) of the sequenced Pyrenean desmans.
**Table S11**. Heterozygosity values in exons of the MHC-I-α and olfactory receptor genes of the sequenced Pyrenean desmans.
**Table S12**. Proportion in ROH regions of exons of the MHC-I-α and olfactory receptor genes of the sequenced Pyrenean desmans.
**Figure S1**. Distributions showing the main features of the Bloom filter-based genome assembly of the Pyrenean desman and the predicted protein-coding genes.
**Figure S2**. GC content of the autosomal scaffolds longer than 10 Mb.
**Figure S3**. Dotplot of the comparison of the dromedary assemblies.
**Figure S4**. Phylogenetic trees of the MHC-I-α and olfactory receptor genes.
**Figure S5**. Historical effective population size inferred from the Pyrenean desman genomes by PSMC with 100 bootstrap replicates.
**Figure S6**. Genome-wide heterozygosity rate for different mammalian species of conservation concern.

**Table S1.** Specimens used in this study with information about their sex, sampling year, locality, geographical area and autonomous commnunity. Two specimens used in a previous study are indicated.

| Specimen code | Sex | Year | Locality | Geographical area |
|---|---|---|---|---|
| IBE-C5619 | Male | 2017 | Torán | Eastern Pyrenees (Catalunya) |
| IBE-C2769 | Male | 1999 | Ezpelura-Urrotz | Western Pyrenees (Navarra) |
| IBE-C3734 (1) | Female | 2011 | Oja | Northwestern Iberian Range (La Rioja) |
| IBE-C3773 (1) | Male | 2011 | Mayor | Southeastern Iberian Range (La Rioja) |
| IBE-BC2778 | Male | 2019 | Hija de Dios | Central System (Castilla y León) |
| IBE-C6507 | Male | 2018 | Requejo | West of the Iberian Peninsula (Castilla y León) |

[1] Escoda L, González-Esteban J, Gómez A, Castresana J (2017) Using relatedness networks to infer contemporary dispersal: application to the endangered mammal *Galemys pyrenaicus*. *Molecular Ecology*, **26**, 3343–3357.

**Table S2.** Summary statistics of the *de novo* genome sequencing data.

| Library name | Library insert size (bp) | Read length (bp) | Raw reads | Filtered reads | Filtered bases | Coverage |
|---|---|---|---|---|---|---|
| C5619_20Gb | 350 | 150 | 168,103,284 | 158,251,220 | 23,737,683,000 | 13.0 |
| C5619_50Gb | 350 | 150 | 353,994,004 | 333,733,964 | 50,060,094,600 | 27.4 |
| C5619_60Gb | 550 | 150 | 663,416,860 | 625,115,962 | 93,767,394,300 | 51.3 |
| C5619_5kb | 5,000 | 150 | 69,131,334 | 69,034,668 | 10,313,274,039 | 5.6 |
| C5619_9kb | 9,000 | 150 | 313,352,690 | 290,403,706 | 43,426,665,509 | 23.8 |
| **Total** | | | **1,567,998,172** | **1,476,539,520** | **221,305,111,448** | **121.0** |

**Table S3.** Summary statistics of genome sequencing data of the additional individuals.

| Library name | Library insert size (bp) | Read length (bp) | Raw reads | Filtered reads | Filtered bases | Coverage |
|---|---|---|---|---|---|---|
| C2769_20Gb | 350 | 150 | 133,311,010 | 125,595,550 | 18,839,332,500 | 10.3 |
| C3734_20Gb | 350 | 150 | 164,266,428 | 156,862,252 | 23,529,337,800 | 12.9 |
| C3773_20Gb | 350 | 150 | 180,798,280 | 171,444,472 | 25,716,670,800 | 14.1 |
| BC2778_50Gb | 350 | 150 | 423,518,938 | 406,106,310 | 60,915,946,500 | 33.3 |
| C6507_20Gb | 350 | 150 | 186,336,338 | 175,033,874 | 26,255,081,100 | 14.4 |

**Table S4.** Summary statistics of the genome assemblies using different combinations of parameters in ABySS. All the assemblies have the following parameters in common: Bloom filter size (B) = 80G, number of Bloom filter hash functions (H) = 4, and minimum untig size required for building contigs (s) = 1000. The final assembly chosen is shown in bold.

| | Abyss parameters | | | | Contigs | | | | | Scaffolds | | | | | BUSCO analysis | | | | |
|---|---|---|---|---|---|---|---|---|---|---|---|---|---|---|---|---|---|---|---|
| # | k | kc | n | N | Number | N50 | Total length (bp) | Largest sequence (bp) | N's | Number | N50 | Total length (bp) | Largest sequence (bp) | N's | Comp. | Sing. | Dup. | Frag. | Miss. |
| 1 | 80 | 2 | 5 | 5 | 111,433 | 32,553 | 1,785,457,123 | 291,897 | 332,861 | 26,042 | 1,200,095 | 1,830,800,637 | 9,588,943 | 47,235,273 | 3,899 | 3,877 | 22 | 133 | 72 |
| 2 | 80 | 2 | 5 | 10 | 111,433 | 32,553 | 1,785,457,123 | 291,897 | 332,861 | 23,940 | 5,951,183 | 1,832,660,941 | 23,566,475 | 49,089,317 | 3,940 | 3,919 | 21 | 103 | 61 |
| 3 | 80 | 2 | 10 | 5 | 113,947 | 31,323 | 1,785,225,099 | 291,897 | 397,393 | 25,437 | 1,269,001 | 1,830,952,436 | 12,846,275 | 47,722,165 | 3,904 | 3,885 | 19 | 127 | 73 |
| 4 | 80 | 2 | 10 | 10 | 113,947 | 31,323 | 1,785,225,099 | 291,897 | 397,393 | 23,437 | 5,697,658 | 1,832,922,782 | 23,203,185 | 49,721,941 | 3,944 | 3,924 | 20 | 97 | 63 |
| 5 | 80 | 3 | 5 | 5 | 109,933 | 33,225 | 1,785,420,032 | 385,381 | 400,687 | 25,857 | 1,213,311 | 1,830,181,266 | 8,465,635 | 46,605,776 | 3,903 | 3,880 | 23 | 128 | 73 |
| 6 | 80 | 3 | 5 | 10 | 109,933 | 33,225 | 1,785,420,032 | 385,381 | 400,687 | 23,766 | 6,073,268 | 1,832,267,957 | 28,195,572 | 48,689,944 | 3,941 | 3,919 | 22 | 99 | 64 |
| 7 | 80 | 3 | 10 | 5 | 112,616 | 31,855 | 1,785,480,117 | 347,451 | 460,921 | 25,328 | 1,249,154 | 1,830,590,970 | 9,531,084 | 47,091,628 | 3,906 | 3,886 | 20 | 129 | 69 |
| 8 | 80 | 3 | 10 | 10 | 112,616 | 31,855 | 1,785,480,117 | 347,451 | 460,921 | 23,302 | 5,554,286 | 1,832,778,480 | 28,196,456 | 49,317,935 | 3,947 | 3,928 | 19 | 94 | 63 |
| 9 | 90 | 2 | 5 | 5 | 82,588 | 45,555 | 1,796,039,717 | 443,520 | 349,179 | 16,098 | 1,791,614 | 1,827,110,659 | 17,686,848 | 32,988,668 | 3,934 | 3,917 | 17 | 100 | 70 |
| 10 | 90 | 2 | 5 | 10 | 82,588 | 45,555 | 1,796,039,717 | 443,520 | 349,179 | 14,975 | 7,181,995 | 1,829,147,288 | 23,672,106 | 35,023,894 | 3,952 | 3,931 | 21 | 85 | 67 |
| 11 | 90 | 2 | 10 | 5 | 85,338 | 43,273 | 1,796,092,999 | 443,520 | 417,713 | 15,683 | 1,807,790 | 1,827,675,698 | 9,450,852 | 33,699,235 | 3,929 | 3,907 | 22 | 106 | 69 |
| 12 | 90 | 2 | 10 | 10 | 85,338 | 43,273 | 1,796,092,999 | 443,520 | 417,713 | 14,622 | 7,459,917 | 1,829,836,990 | 36,183,241 | 35,866,583 | 3,947 | 3,925 | 22 | 90 | 67 |
| 13 | 90 | 3 | 5 | 5 | 82,394 | 46,348 | 1,795,401,409 | 410,797 | 458,625 | 16,825 | 1,722,795 | 1,826,834,836 | 9,026,015 | 33,376,920 | 3,918 | 3,899 | 19 | 111 | 75 |
| 14 | 90 | 3 | 5 | 10 | 82,394 | 46,348 | 1,795,401,409 | 410,797 | 458,625 | 17,247 | 7,154,225 | 1,828,635,784 | 34,866,347 | 35,215,360 | 3,953 | 3,932 | 21 | 88 | 63 |
| 15 | 90 | 3 | 10 | 5 | 85,015 | 44,011 | 1,795,415,076 | 473,012 | 529,804 | 18,377 | 1,755,848 | 1,827,249,887 | 11,938,933 | 33,967,457 | 3,919 | 3,902 | 17 | 116 | 69 |
| 16 | 90 | 3 | 10 | 10 | 85,015 | 44,011 | 1,795,415,076 | 473,012 | 529,804 | 15,002 | 7,263,463 | 1,829,516,963 | 36,159,321 | 36,235,826 | 3,948 | 3,930 | 18 | 89 | 67 |
| 17 | 100 | 2 | 5 | 5 | 63,049 | 63,603 | 1,803,404,661 | 480,138 | 442,655 | 13,929 | 2,363,627 | 1,826,436,552 | 17,794,570 | 25,161,174 | 3,936 | 3,914 | 22 | 97 | 71 |
| 18 | 100 | 2 | 5 | 10 | 63,049 | 63,603 | 1,803,404,661 | 480,138 | 442,655 | 12,831 | 8,511,288 | 1,828,107,683 | 34,981,249 | 26,855,549 | 3,947 | 3,922 | 25 | 92 | 65 |
| 19 | 100 | 2 | 10 | 5 | 64,927 | 60,334 | 1,803,400,543 | 463,860 | 501,639 | 14,683 | 2,457,278 | 1,826,595,599 | 12,564,118 | 25,472,454 | 3,940 | 3,916 | 24 | 97 | 67 |
| **20** | **100** | **2** | **10** | **10** | **64,927** | **60,334** | **1,803,400,543** | **463,860** | **501,639** | **12,306** | **8,503,682** | **1,828,347,170** | **36,404,611** | **27,224,426** | **3,953** | **3,931** | **22** | **86** | **65** |
| 21 | 100 | 3 | 5 | 5 | 69,357 | 63,700 | 1,800,943,157 | 726,128 | 648,716 | 18,162 | 2,158,393 | 1,828,462,988 | 12,351,970 | 29,828,976 | 3,936 | 3,914 | 22 | 97 | 71 |
| 22 | 100 | 3 | 5 | 10 | 69,357 | 63,700 | 1,800,943,157 | 726,128 | 648,716 | 17,270 | 7,464,961 | 1,830,000,858 | 34,975,378 | 31,398,726 | 3,946 | 3,923 | 23 | 90 | 68 |
| 23 | 100 | 3 | 10 | 5 | 70,076 | 60,387 | 1,801,259,151 | 713,669 | 747,718 | 16,607 | 2,285,473 | 1,828,365,895 | 10,356,824 | 29,475,535 | 3,934 | 3,913 | 21 | 103 | 67 |
| 24 | 100 | 3 | 10 | 10 | 70,076 | 60,387 | 1,801,259,151 | 713,669 | 747,718 | 15,878 | 7,378,031 | 1,830,046,073 | 35,735,109 | 31,189,660 | 3,948 | 3,925 | 23 | 93 | 63 |
| 25 | 110 | 2 | 5 | 5 | 64,540 | 76,058 | 1,806,041,319 | 686,611 | 650,423 | 20,494 | 2,922,818 | 1,830,260,978 | 14,191,065 | 26,895,880 | 3,938 | 3,912 | 26 | 102 | 64 |
| 26 | 110 | 2 | 5 | 10 | 64,540 | 76,058 | 1,806,041,319 | 686,611 | 650,423 | 18,701 | 9,154,820 | 1,832,069,487 | 31,306,718 | 28,726,537 | 3,950 | 3,925 | 25 | 90 | 64 |
| 27 | 110 | 2 | 10 | 5 | 63,849 | 73,186 | 1,806,339,970 | 686,611 | 742,276 | 17,004 | 3,310,031 | 1,830,799,709 | 17,730,278 | 27,184,354 | 3,937 | 3,913 | 24 | 100 | 67 |
| 28 | 110 | 2 | 10 | 10 | 63,849 | 73,186 | 1,806,339,970 | 686,611 | 742,276 | 16,524 | 9,577,427 | 1,831,694,757 | 31,308,620 | 28,141,420 | 3,956 | 3,934 | 22 | 82 | 66 |
| 29 | 110 | 3 | 5 | 5 | 99,452 | 70,759 | 1,793,541,866 | 710,746 | 1,036,511 | 47,086 | 2,620,137 | 1,838,978,192 | 11,612,014 | 48,562,092 | 3,892 | 3,868 | 24 | 133 | 79 |
| 30 | 110 | 3 | 5 | 10 | 99,452 | 70,759 | 1,793,541,866 | 710,746 | 1,036,511 | 48,094 | 9,109,270 | 1,835,154,356 | 36,456,462 | 44,722,154 | 3,900 | 3,876 | 24 | 117 | 87 |
| 31 | 110 | 3 | 10 | 5 | 95,217 | 68,549 | 1,794,502,044 | 686,728 | 1,196,453 | 41,311 | 2,648,512 | 1,838,754,245 | 13,241,708 | 47,426,515 | 3,892 | 3,871 | 21 | 128 | 84 |
| 32 | 110 | 3 | 10 | 10 | 95,217 | 68,549 | 1,794,502,044 | 686,728 | 1,196,453 | 42,202 | 8,942,909 | 1,836,068,300 | 26,818,792 | 44,770,810 | 3,926 | 3,904 | 22 | 95 | 83 |

k: size of k-mer; kc: minimum k-mer count threshold for Bloom filter assembly; n: minimum number of pairs required for building contigs; N: minimum number of pairs required for building scaffolds; Comp.: Complete BUSCOs; Sing.: Complete single-copy BUSCOs; Dup.: Complete duplicated BUSCOs; Frag.: Fragmented BUSCOs, Miss.: Missing BUSCOs.

**Table S5.** Summary statistics of the genome assembly. All the statistics are based on contigs of size >= 500 bp.

| **Statistics** | **Contigs** | **Scaffolds** |
|---|---:|---:|
| Sequence count | 64,927 | 12,306 |
| Total length (bp) | 1,803,400,543 | 1,828,347,170 |
| Largest contig (bp) | 463,860 | 36,404,611 |
| N's | 501,639 | 27,224,426 |
| N50 (count) | 60,334 (8,720) | 8,503,682 (66) |
| N75 (count) | 31,301 (19,089) | 3,815,754 (140) |

**Table S6.** Summary of the BUSCO analysis using 4,104 mammalian single-copy orthologs database.

| Statistics | Count | Ratio (%) |
|---|---|---|
| Complete BUSCOs | 3,953 | 96.3 |
| Complete single-copy BUSCOs | 3,931 | 95.8 |
| Complete duplicated BUSCOs | 22 | 0.5 |
| Fragmented BUSCOs | 86 | 2.1 |
| Missing BUSCOs | 65 | 1.6 |

**Table S7.** Summary statistics of the genome assemblies of the dromedary using ABySS. The standard assembly was publised in Fitak et al. (2016)[1]. The Bloom-filter assembly was produced using the raw sequence data from Fitak et al. (2016)[1] and the best Bloom filter options found for the desman.

| | Scaffolds | | | | | BUSCO analysis | | | | |
|---|---|---|---|---|---|---|---|---|---|---|
| **Assembly** | **Number** | **N50** | **Largest sequence (bp)** | **N's** | **Total length (bp)** | **Comp.** | **Sing.** | **Dup.** | **Frag.** | **Miss.** |
| Standard | 35,752 | 1,482,444 | 9,719,801 | 53,439,631 | 2,055,063,633 | 3,909 | 3,886 | 23 | 112 | 83 |
| Bloom filter | 45,507 | 2,086,244 | 13,669,667 | 74,648,979 | 1,999,645,312 | 3,852 | 3,835 | 17 | 142 | 110 |

Comp.: Complete BUSCOs; Sing.: Complete Single-Copy BUSCOs; Dup.: Complete Duplicated BUSCOs; Frag.: Fragmented BUSCOs, Miss.: Missing BUSCOs.

[1] Fitak, R. R., Mohandesan, E., Corander, J., & Burger, P. A. (2016). The *de novo* genome assembly and annotation of a female domestic dromedary of North African origin. Molecular Ecology Resources, 16 (1), 314–324.

**Table S8.** Summary statistics of the repetitive elements identified with RepeatMasker.

| **TE class** | **Count** | **Length (bp)** | **Ratio (%)** |
|---|---|---|---|
| SINEs: | 746,460 | 150,640,913 | 8.24 |
| Alu/B1 | 28 | 788 | 0.00 |
| MIRs | 266,561 | 35,470,149 | 1.94 |
| LINEs: | 548,054 | 204,676,726 | 11.19 |
| LINE1 | 361,334 | 161,962,297 | 8.86 |
| LINE2 | 157,413 | 37,140,022 | 2.03 |
| L3/CR1 | 23,967 | 4,506,547 | 0.25 |
| RTE | 4,583 | 968,770 | 0.05 |
| LTR elements: | 264,663 | 80,383,117 | 4.40 |
| ERVL | 51,842 | 18,704,034 | 1.02 |
| ERVL-MaLRs | 82,131 | 24,678,381 | 1.35 |
| ERV_classI | 87,954 | 29,637,393 | 1.62 |
| ERV_classII | 24,854 | 2,370,471 | 0.13 |
| DNA elements: | 199,476 | 38,364,850 | 2.10 |
| hAT-Charlie | 108,626 | 19,788,936 | 1.08 |
| TcMar-Tigger | 41,077 | 9,333,173 | 0.51 |
| Unclassified: | 3,288 | 570,873 | 0.03 |
| Small RNA: | 20,790 | 1,797,159 | 0.10 |
| Satellites: | 55,522 | 19,797,132 | 1.08 |
| Simple repeats: | 405,554 | 18,609,900 | 1.02 |
| Low complexity: | 84,860 | 4,446,792 | 0.24 |
| **Total interspersed repeats** | | **474,636,479** | **25.96** |

**Table S9.** Autosomal genome-wide heterozygosity of the sequenced Pyrenean desmans, given in SNPs/Mb, for two different values of minimum depth of coverage. Averages for all individuals and for the three individuals with ddRAD data are also given.

| | Minimum depth ≥ 12 | | | Minimum depth ≥ 5 | | | |
|---|---|---|---|---|---|---|---|
| **Specimen code** | **Heterozygous positions** | **Total positions** | **Genomic heterozygosity** | **Heterozygous positions** | **Total positions** | **Genomic heterozygosity** | **ddRAD heterozygosity** |
| IBE-C5619 | 20,649 | 1,695,877,553 | 12 | 20,996 | 1,697,087,555 | 12 | - |
| IBE-C2769 | 32,231 | 266,881,822 | 121 | 155,806 | 1,543,525,767 | 101 | - |
| IBE-C3734 | 163,835 | 629,146,242 | 260 | 384,285 | 1,643,656,575 | 234 | 179 |
| IBE-C3773 | 193,673 | 806,670,037 | 240 | 371,534 | 1,663,708,855 | 223 | 169 |
| IBE-BC2778 | 195,602 | 1,680,320,379 | 116 | 199,603 | 1,692,398,487 | 118 | - |
| IBE-C6507 | 413,089 | 892,035,963 | 463 | 741,672 | 1,668,804,108 | 444 | 359 |
| **Average (All)** | | | 202 | | | 189 | |
| **Average (3 ind.)** | | | 321 | | | 300 | 236 |

**Table S10.** Runs of homozygosity (ROH) of the sequenced Pyrenean desmans calculated with different methods in comparison with the inbreeding coefficients calculated from ddRAD data for three individulas. Averages for all individuals and for the three individuals with ddRAD data are also given.

| **Specimen code** | **PLINK** | **BCFtools/ RoH** | **ROHan** | **Proportion of homozygous 100-kb windows** | **ddRAD inbreeding coefficient** |
|---|---|---|---|---|---|
| IBE-C5619 | 0.97 | 0.83 | 0.53 | 0.69 | - |
| IBE-C2769 | 0.79 | 0.57 | 0.62 | 0.47 | - |
| IBE-C3734 | 0.55 | 0.36 | 0.39 | 0.35 | 0.34 |
| IBE-C3773 | 0.57 | 0.36 | 0.39 | 0.34 | 0.40 |
| IBE-BC2778 | 0.74 | 0.51 | 0.54 | 0.49 | - |
| IBE-C6507 | 0.19 | 0.09 | 0.11 | 0.10 | 0.09 |
| **Average (All)** | **0.64** | **0.45** | **0.43** | **0.41** | |
| **Average (3 ind.)** | 0.44 | 0.27 | 0.30 | 0.26 | 0.28 |

**Table S11.** Heterozygosity values in exons of the MHC-I-α and olfactory receptor genes of the sequenced Pyrenean desmans, given in SNPs/Mb, in comparison with the expected heterozigosity (heterozigosity in the whole genome with minimum depth of coverage ≥ 5 extrated from Table S9). The heterozigosity excess with respect to the expected proportion is given in brackets. Average across individuals as well as the number of total exon positions analyzed for each protein class are also given (the analysis is based on autosomal scaffolds > 40,000 bp).

| **Specimen code** | **Expected heterozygosity** | **All exons** | **MHC-I-α** | **Olfactory receptor** |
|---|---|---|---|---|
| IBE-C5619 | 12 | 23 (1.9x) | 114 (9.5x) | 53 (4.4x) |
| IBE-C2769 | 101 | 91 (0.9x) | 114 (1.1x) | 147 (1.5x) |
| IBE-C3734 | 234 | 217 (0.9x) | 2,235 (9.6x) | 632 (2.7x) |
| IBE-C3773 | 223 | 198 (0.9x) | 1,932 (8.7x) | 941 (4.2x) |
| IBE-BC2778 | 118 | 129 (1.1x) | 417 (3.5x) | 809 (6.9x) |
| IBE-C6507 | 444 | 368 (0.8x) | 7,500 (16.9x) | 1,050 (2.4x) |
| **Average** | **189** | **171 (0.9x)** | **2,052 (10.9x)** | **605 (3.2x)** |
| Exon positions analyzed | | 33,533,344 | 26,400 | 469,725 |

**Table S12.** Proportion in ROH regions of exons of the MHC-I-α and olfactory receptor genes of the sequenced Pyrenean desmans in comparison with the expected proportion (proportion of homozygous 100-kb windows taken from Table S10). The p-value is given in brackets. Average across individuals as well as the total number of exons analyzed for each protein class are also given (the analysis is based on autosomal scaffolds > 100,000 bp).

| Specimen code | Expected proportion | All exons | MHC-I-α | Olfactory receptor |
|---|---|---|---|---|
| IBE-C5619 | 0.69 | 0.62 (0.00) | 0.11 (0.00) | 0.53 (0.00) |
| IBE-C2769 | 0.47 | 0.44 (0.00) | 0.48 (0.50) | 0.32 (0.00) |
| IBE-C3734 | 0.35 | 0.31 (0.00) | 0.00 (0.00) | 0.25 (0.02) |
| IBE-C3773 | 0.34 | 0.31 (0.00) | 0.00 (0.00) | 0.28 (0.12) |
| IBE-BC2778 | 0.49 | 0.46 (0.00) | 0.15 (0.02) | 0.28 (0.00) |
| IBE-C6507 | 0.10 | 0.09 (0.00) | 0.00 (0.00) | 0.00 (0.00) |
| **Average** | **0.41** | **0.37** | **0.12** | **0.28** |
| Exons analyzed | | 173,390 | 115 | 479 |

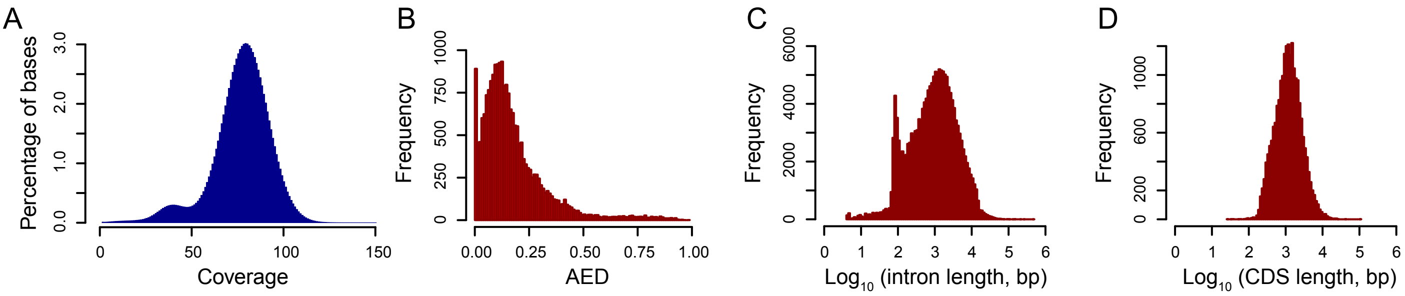


**Figure S1.** Distributions showing the main features of the Bloom filter-based genome assembly of the Pyrenean desman and the predicted protein-coding genes. (A) Coverage of the short-insert sequencing data. (B) Annotation edit distances (AED) of the predicted genes. (C) Logarithm of intron length of the predicted genes. (D) Logarithm of coding sequence (CDS) length of the predicted genes.

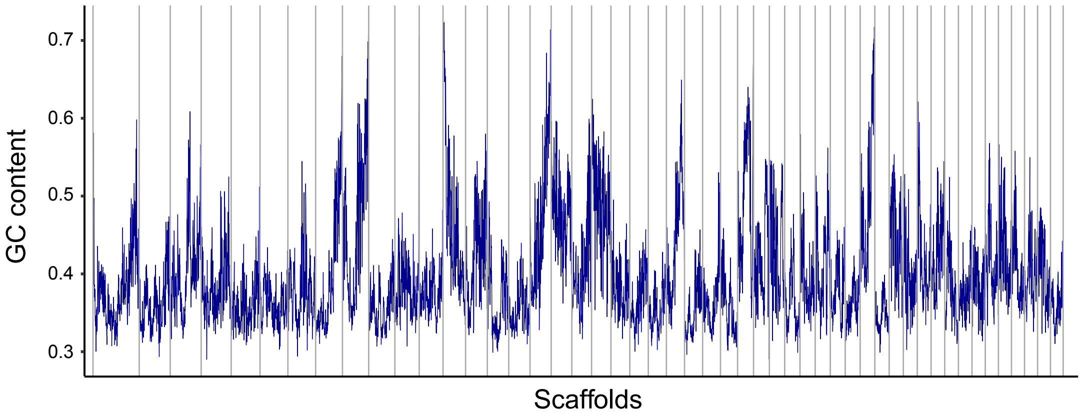


**Figure S2.** GC content of the autosomal scaffolds longer than 10 Mb.

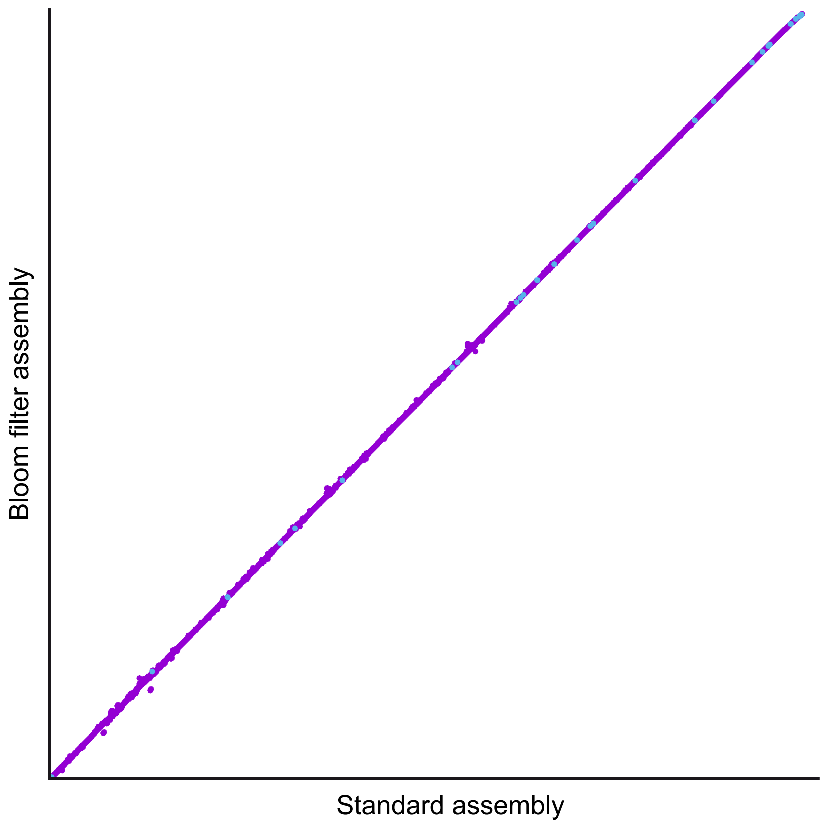


**Figure S3.** Dotplot of the comparison of dromedary assemblies generated with MUMmer. Forward matches are shown in purple and reverse matches in blue.

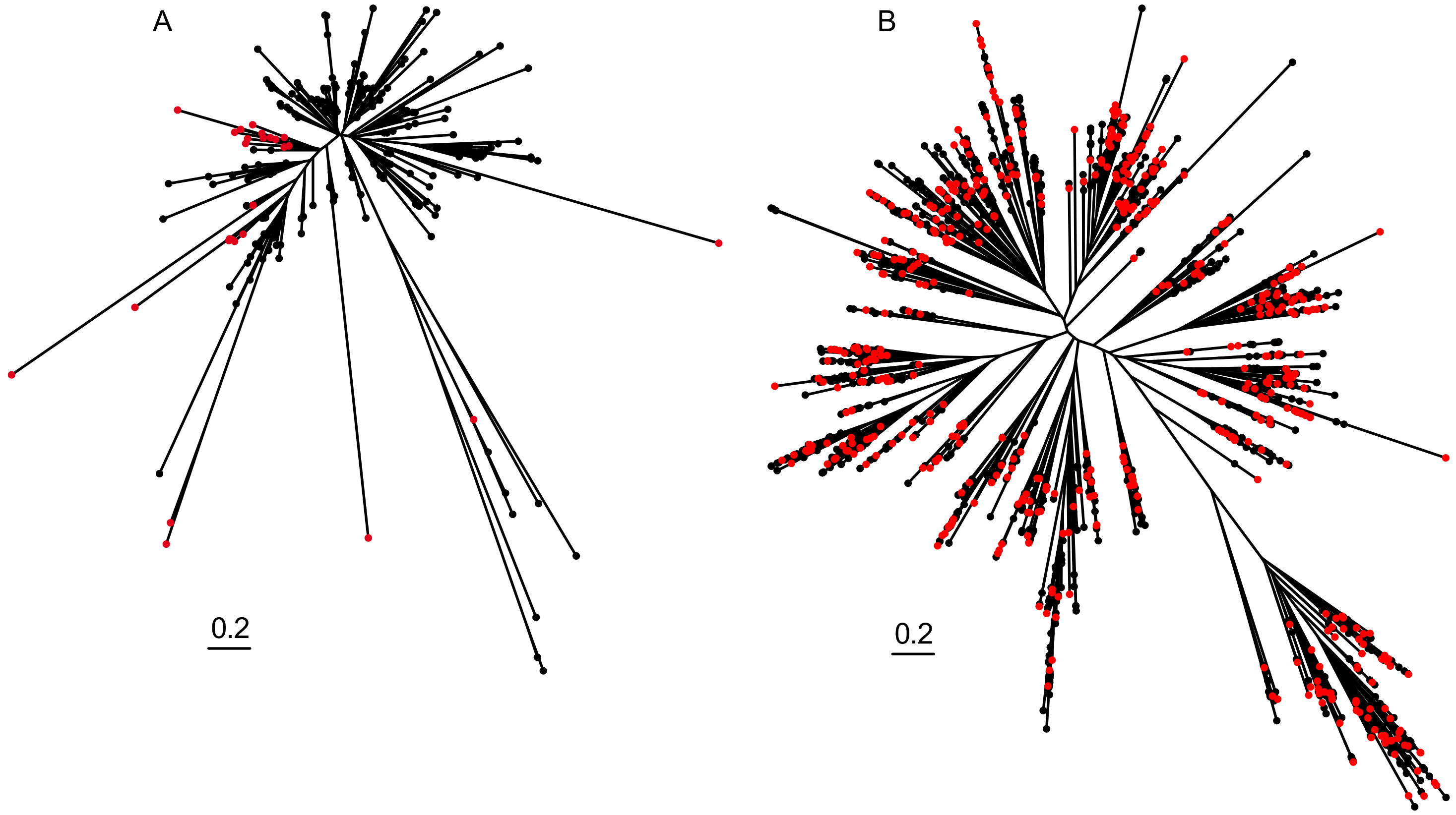


**Figure S4.** Maximum-likelihood phylogenetic trees of 26 Pyrenean desman MHC-I-α genes together with those of the mole *Condylura cristata*, human and several mammalas from Abduriyim et al. 2019 (A), and 529 Pyrenean desman olfactory receptor genes of Pyrenean desman together with those of *Condylura cristata* and human (B), constructed from the amino acid alignments. The Pyrenean desman sequences are shown with a red circle and those of other mammals with a black circle. The scale represents 0.2 substitutions per position and is the same in both cases.

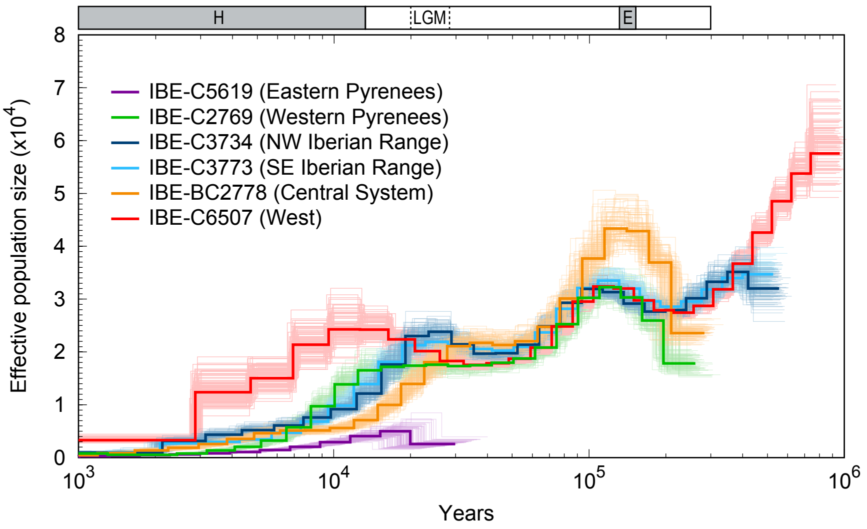


**Figure S5.** Historical effective population size inferred from the Pyrenean desman genomes by PSMC. The lighter coloured lines of the same colour represent the 100 bootstrap replicates. The result is scaled with a mutation rate ($\mu$) of 5 x $10^{-9}$ mutations/site/generation and an average generation time of 2 years. The last two interglacial periods, Holocene (H) and Eemian (E), are indicated with grey boxes and the Last Glacial Maximum (LGM) with dashed lines.

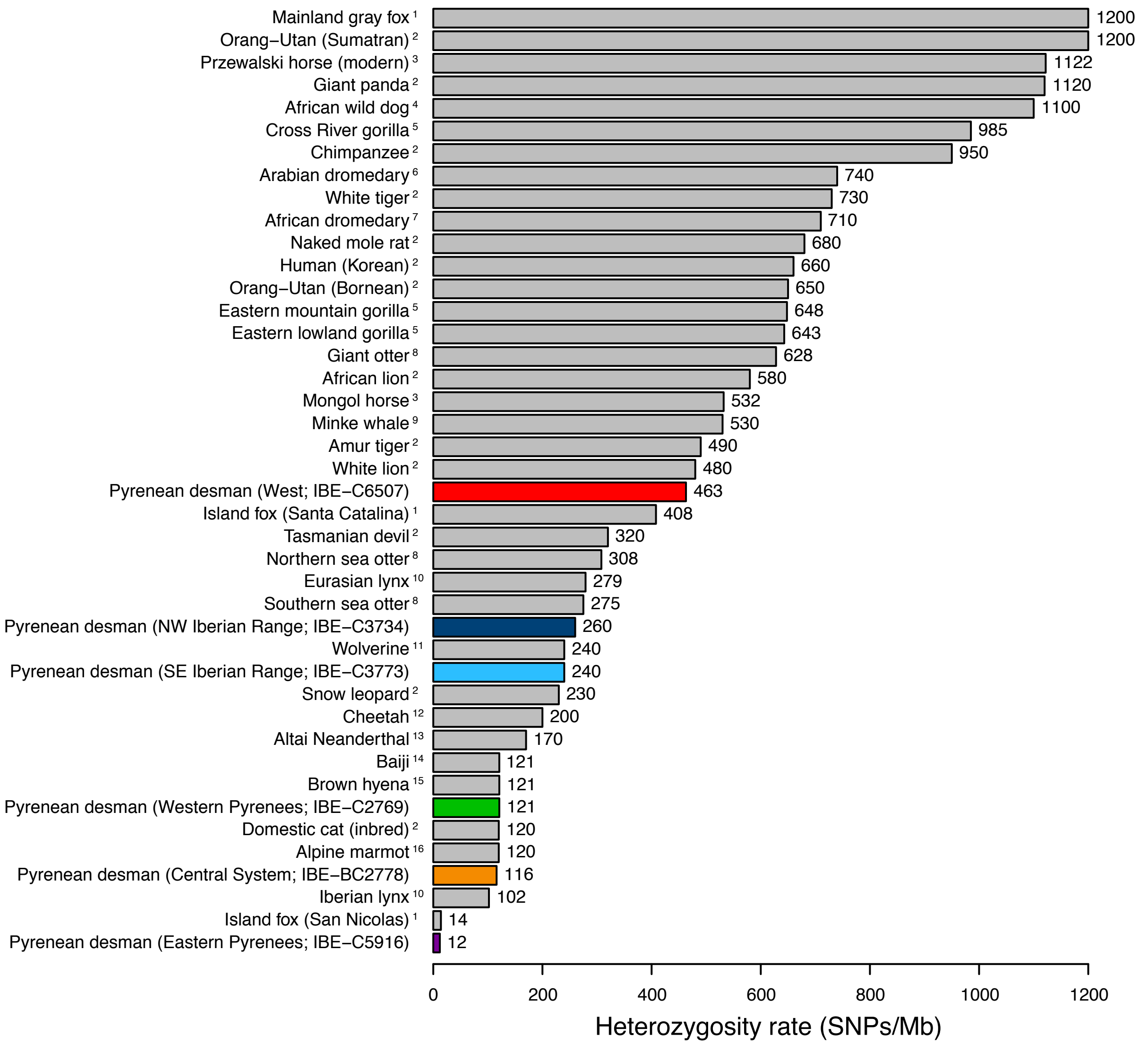



[1] Robinson, J. A. et al. Genomic Flatlining in the Endangered Island Fox. Curr. Biol. 26, 1183–1189 (2016).

[2] Cho, Y. S. et al. The tiger genome and comparative analysis with lion and snow leopard genomes. Nat. Commun. 4, (2013).

[3] Der Sarkissian, C. et al. Evolutionary genomics and conservation of the endangered Przewalski's horse. Curr. Biol. 25, 2577–2583 (2015).

[4] Armstrong, E. E. et al. Cost-effective assembly of the African wild dog ( Lycaon pictus ) genome using linked reads. Gigascience 8, 1–10 (2019).

[5] Xue, Y. et al. Mountain gorilla genomes reveal the impact of long-term population decline and inbreeding. Science (80-. ). 348, 242–245 (2015).

[6] Wu, H. et al. Camelid genomes reveal evolution and adaptation to desert environments. Nat. Commun. 5, (2014).

[7] Fitak, R. R., Mohandesan, E., Corander, J. & Burger, P. A. The de novo genome assembly and annotation of a female domestic dromedary of North African origin. Mol. Ecol. Resour. 16, 314–324 (2016).

[8] Beichman, A. C. et al. Aquatic Adaptation and Depleted Diversity: A Deep Dive into the Genomes of the Sea Otter and Giant Otter. Mol. Biol. Evol. 36, 2631–2655 (2019).

[9] Yim, H. S. et al. Minke whale genome and aquatic adaptation in cetaceans. Nat. Genet. 46, 88–92 (2014).

[10] Abascal, F. et al. Extreme genomic erosion after recurrent demographic bottlenecks in the highly endangered Iberian lynx. Genome Biol. 17, 251 (2016).

[11] Ekblom, R. et al. Genome sequencing and conservation genomics in the Scandinavian wolverine population. Conserv. Biol. 32, 1301–1312 (2018).

[12] Dobrynin, P. et al. Genomic legacy of the African cheetah, Acinonyx jubatus. Genome Biol. 16, 1–19 (2015).

[13] Prüfer, K. et al. The complete genome sequence of a Neanderthal from the Altai Mountains. Nature 505, 43–49 (2014).

[14] Zhou, X. et al. Baiji genomes reveal low genetic variability and new insights into secondary aquatic adaptations. Nat. Commun. 4, 1–6 (2013).

[15] Westbury, M. V. et al. Extended and continuous decline in effective population size results in low genomic diversity in the world's rarest hyena species, the brown Hyena. Mol. Biol. Evol. 35, 1225–1237 (2018).

[16] Gossmann, T. I. et al. Ice-Age Climate Adaptations Trap the Alpine Marmot in a State of Low Genetic Diversity. Curr. Biol. 29, 1712–1720.e7 (2019).


**Figure S6.** Genome-wide heterozygosity rate for different mammalian species, most of them of conservation concern. Values of different Pyrenean desmans sequenced in this work are shown in color.
